# Simultaneous detection of invasive signal crayfish, endangered white-clawed crayfish and the crayfish plague pathogen using environmental DNA

**DOI:** 10.1101/291856

**Authors:** Chloe Victoria Robinson, Tamsyn M. Uren Webster, Joanne Cable, Joanna James, Sofia Consuegra

## Abstract

Aquatic Invasive Species (AIS) are important vectors for the introduction of novel pathogens which can, in turn, become drivers of rapid ecological and evolutionary change, compromising the persistence of native species. Conservation strategies rely on accurate information regarding presence and distribution of AIS and their associated pathogens to prevent or mitigate negative impacts, such as predation, displacement or competition with native species for food, space or breeding sites. Environmental DNA is increasingly used as a conservation tool for early detection and monitoring of AIS. We used a novel eDNA high-resolution melt curve (HRM) approach to simultaneously detect the UK endangered native crayfish (*Austropotamobius pallipes*), the highly invasive signal crayfish (*Pacifastacus leniusculus*) and their dominant pathogen, *Aphanomyces astaci*, (causative agent of crayfish plague). We validated the approach with laboratory and field samples in areas with known presence or absence of both crayfish species as well as the pathogen, prior to the monitoring of areas where their presence was unknown. We identified the presence of infected signal crayfish further upstream than previously detected in an area where previous intensive eradication attempts had taken place, and the coexistence of both species in plague free catchments. We also detected the endangered native crayfish in an area where trapping had failed. With this method, we could estimate the distribution of native and invasive crayfish and their infection status in a rapid, cost effective and highly sensitive way, providing essential information for the development of conservation strategies in catchments with populations of endangered native crayfish.

## INTRODUCTION

Invasive non-native species have become important drivers of global environmental change (Vitousek et al. 1996), although the importance of their impacts on biodiversity remains controversial (Russell and Blackburn 2017). Their spread has been favoured by human-mediated activities (Crowl et al. 2008) in addition to natural dispersal, and, as a consequence have also become common vehicles for the introduction of novel pathogens (Randolph and Rogers 2010). Invasive non-native species extend the geographic range of the pathogens they carry and facilitate host-switching (Peeler et al. 2011). In turn, pathogens play an important role in the evolution of communities but can also threaten the survival of native populations (Altizer et al. 2003). Co-introductions of parasites with non-native hosts are common; invasive species may bring novel infectious diseases that can infect native competitors, but can also act as hosts and effective dispersers for native diseases (Strauss et al. 2012). Invasive pathogens can have devastating effects on vulnerable native hosts, as their virulence tends to be higher than in the non-native species (Lymbery et al. 2014). Such pathogens seem particularly frequent in freshwater species, potentially reflecting the high susceptibility of freshwater ecosystems to non-native invasions (Moorhouse and Macdonald 2015). Thus, early detection of both non-native hosts and parasites is critical for the control and management of the impacts caused by introduced diseases.

Detection of non-native species often occurs when populations have already established, spread from original source and altered the local environment (Vander Zanden et al. 2010; Zaiko et al. 2014). This is particularly the case in aquatic environments, where juveniles or larvae at the initial stages of introduction often have a patchy distribution, are difficult to identify using morphological techniques, and are easily missed by monitoring programmes (Pochon et al. 2013). Early detection is needed to make management actions such as eradication and control of invasive species more efficient and/or effective (Lodge et al. 2016) and as such is becoming fundamental for the management and control of aquatic invasive species (AIS; Vander Zanden et al. 2010). Analysis of environmental DNA (eDNA), i.e. free DNA molecules released from sources such as faeces, skin, urine, blood or secretions of organisms, is proving increasingly useful for detecting species that are difficult to identify and locate by more traditional and time-consuming methods (Biggs et al. 2015), such as endangered species (Dejean et al. 2011) and AIS at the early stages of their introduction (Bohmann et al. 2014; Dejean et al. 2012). Although still a relatively new tool, eDNA is becoming widely used for conservation (Biggs et al. 2015; Laramie et al. 2015; Spear et al. 2015; Thomsen and Willerslev 2015) and protocols are being refined to increase its accuracy and reliability (Goldberg et al. 2016; Wilson et al. 2016). Quantitative PCR (qPCR) is commonly used to target particular species in eDNA samples (e.g. (Ficetola et al. 2008; Thomsen et al. 2012) and, coupled with *in vitro* controls and amplicon sequencing, has proved a reliable method for the detection of invasive and endangered aquatic species (Klymus et al. 2015; Spear et al. 2015). In addition, qPCR is widely used to detect infectious agents in environmental samples (Guy et al. 2003), and can be particularly useful for the early detection of aquatic pathogens which can be introduced simultaneously with non-native species (Ganoza et al. 2006; Strand et al. 2014). High Resolution Melting (HRM) analysis is a qPCR-based method which facilitates identification of small variations in nucleic acid sequences by differences in the melting temperature of double stranded DNA depending on fragment length and sequence composition (Ririe et al. 1997). Analysis of HRM curves has been widely used for SNP genotyping as a fast method to discriminate species (Yang et al. 2009), including natives and invasives (Ramón-Laca et al. 2014). HRM has the potential for being used in AIS identification, including aquatic invasive pathogens, but it has not yet been applied to their detection from eDNA samples. We have used this method to investigate the distribution of the invasive signal crayfish (*Pacifastacus leniusculus*), carrier of the crayfish plague agent (*Aphanomyces astaci*) which is highly infective for native species (e.g. *Austropotamobius pallipes*), and the potential coexistence between native and invasive crayfish in UK populations.

Invasive non-native crayfish have been globally introduced, mainly for human consumption, and are known to seriously impact native ecosystems through predation, competition, disease transmission and hybridisation (e.g. Lodge et al. 2012). In Europe, non-indigenous crayfish mostly of North American origin have outnumbered their native counterparts in much of their range and represent one of the main threats to their persistence (Holdich et al. 2009). The distribution and abundance of native European crayfish species has been strongly influenced by high mortality rates associated with contracting crayfish plague (Schrimpf et al. 2012) through the introduction of North American freshwater crayfish around 1850 (Alderman 1996). *P. leniusculus* was one of the first non-native species introduced to Europe and in the UK is displacing the native crayfish (*A. pallipes*) which has been classified as endangered in the UK (IUCN 2017). Its success has been attributed to preadaptation, niche plasticity, the aggressive nature of the species (Chapple et al. 2012; Pintor et al. 2008) and/or the competitive advantage provided by the crayfish plague (Bubb, Thom, and Lucas 2006; Dunn et al. 2012; Edgerton et al. 2004; Griffiths, Collen and Armstrong, 2004).

By using a novel approach to simultaneously identify both AIS and their major associated pathogens, we analysed the distribution of the highly invasive signal crayfish (*P. leniusculus*), the native crayfish (*A. pallipes*) and the crayfish plague pathogen (*A. astaci*) in areas where the presence of the signal crayfish is severely impacting the native populations, to identify potential areas of coexistence and refugia for the native species. We expected to find coexisting populations of both species more likely in locations where the crayfish plague has been historically and continually absent.

## MATERIALS AND METHODS

### *EX SITU* OPTIMISATION OF eDNA METHODS

In order to optimise eDNA protocols an *ex-situ* pilot experiment was conducted by placing individual *P. leniusculus* in three isolated tanks, each with 2 L of water. After 24 hours, they were removed and two 15 mL water samples were taken from each tank. The sampling was repeated 24 and 48 hours after removal. Two ultrapure water blanks and four tank blanks (with no crayfish in) were also taken as controls during each sampling period. Immediately after collection, a standard method of preserving and extracting eDNA was applied by the addition of 33 mL of absolute ethanol and 1.5 mL of 3M sodium acetate to samples and subsequent storage at -20°C for a minimum of 24 hours before DNA extraction (Ficetola et al. 2008). To recover precipitated DNA, samples were centrifuged to create a DNA pellet. The supernatant was discarded and the remaining pellet was air-dried before being subjected to DNA extraction. Extraction blanks consisting of ultrapure water in place of sampled water and tank blanks were used to test for any cross-contamination of the samples. Similarly, nine 15 mL water samples were taken, along with a system blank, at a local hatchery containing a population of *A. pallipes*, to test detection levels of native crayfish in aqueous eDNA samples.

### STUDY POPULATIONS AND eDNA SAMPLE COLLECTION

We sampled six locations in the River Wye catchment and seven additional sites in the River Taff catchment, both in Wales, UK (Figure 1a-c), as well as a total of 29 sites in two catchments from Southern England, the Itchen and Medway rivers (Figure 1c; Table 1), all of them introduced c.1970. Records of the introduction of signal crayfish in Europe are very limited, but some evidence suggests that between 1976 and 1978 around 150,000 juvenile signal crayfish were introduced into Britain and other European countries from a hatchery in Simontorp, Sweden, which originally imported them from Lake Tahoe in California and Nevada, USA, in 1969 (Holdich and Lowery, 1988). After the Simontorp introductions, crayfish began to be imported directly from different American hatcheries (Holdich and Lowery, 1988), suggesting that the current populations could have different origins, and potentially initial infection status.

**Figure 1a.**
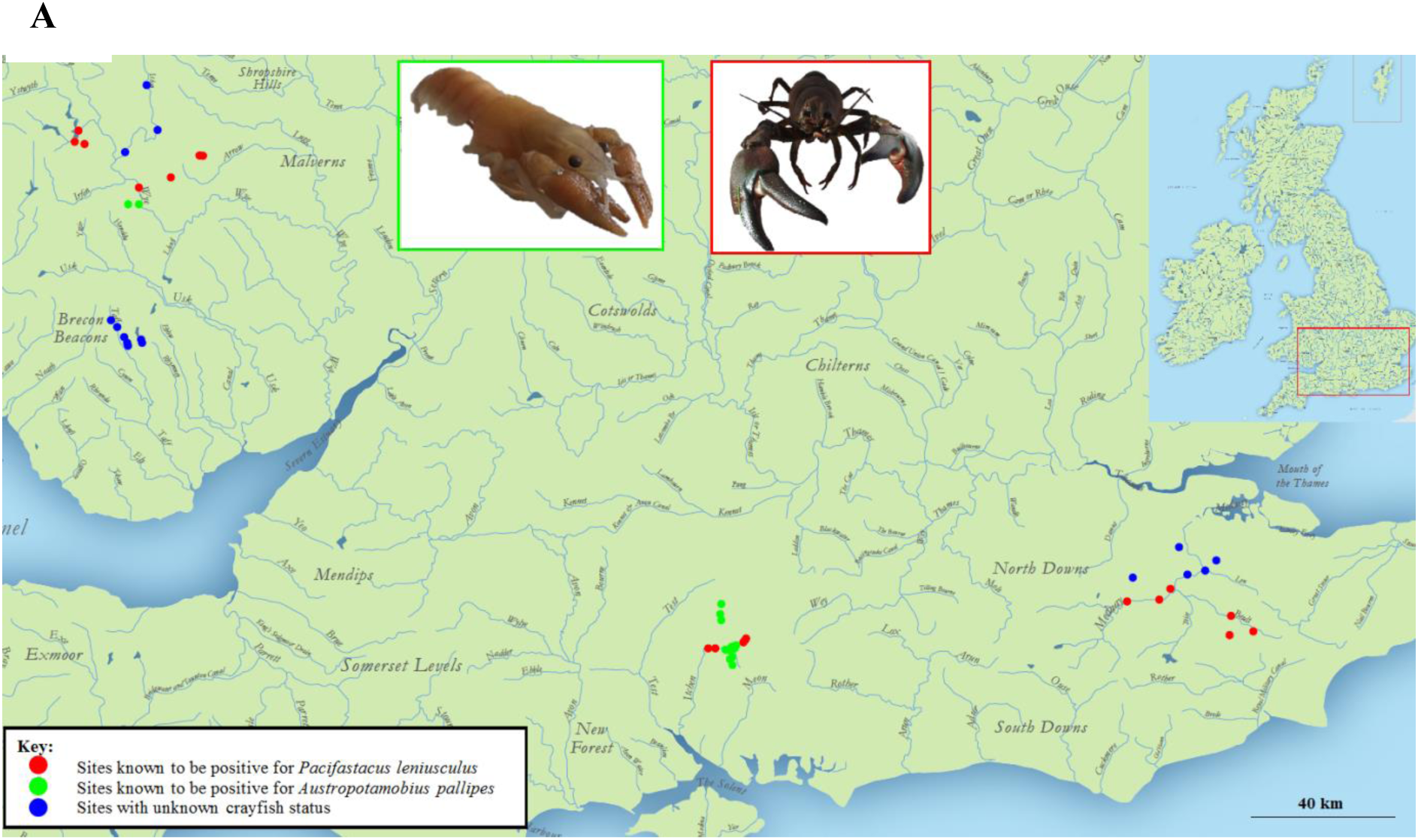
eDNA sampling sites for England (Medway and Itchen) and Wales (Wye and Taff) in tributaries with known presence of *Pacifastacus leniusculus* individuals (red circle), *Austropotamobius pallipes* (green circle) or without information regarding crayfish status (blue circle). Each point represents a locality where between three and nine water samples were collected. (*Austropotamobius pallipes* photograph ©Chloe Robinson; *Pacifastacus leniusculus* photograph ©Rhidian Thomas).

**Figure 1b.**
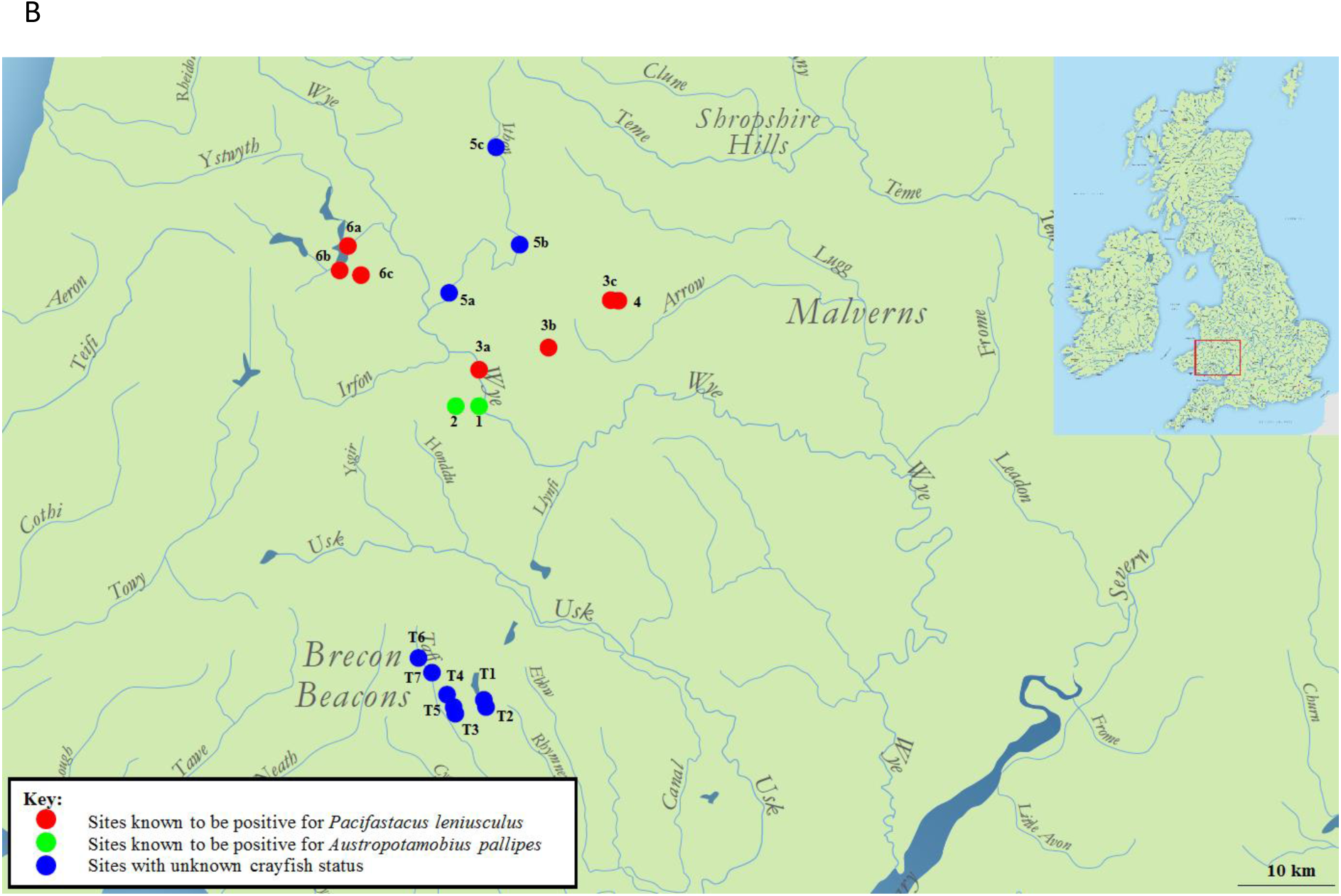
Location of the rivers Wye and Taff eDNA sampling sites in Wales. Wye sites 1 and 2 (Sgithwen Brook) were confirmed for crayfish species *Austropotamobius pallipes*; sites 3 (Bachowey), 4 (Bachowey) and 6 (Duhonw) were confirmed for crayfish species *Pacifastacus leniusculus* and site 5 (Edw) had unknown crayfish status. Taff sites T1 to T7 all had unknown crayfish presence status. Each point corresponds to between three and nine water samples collected

**Figure 1c.**
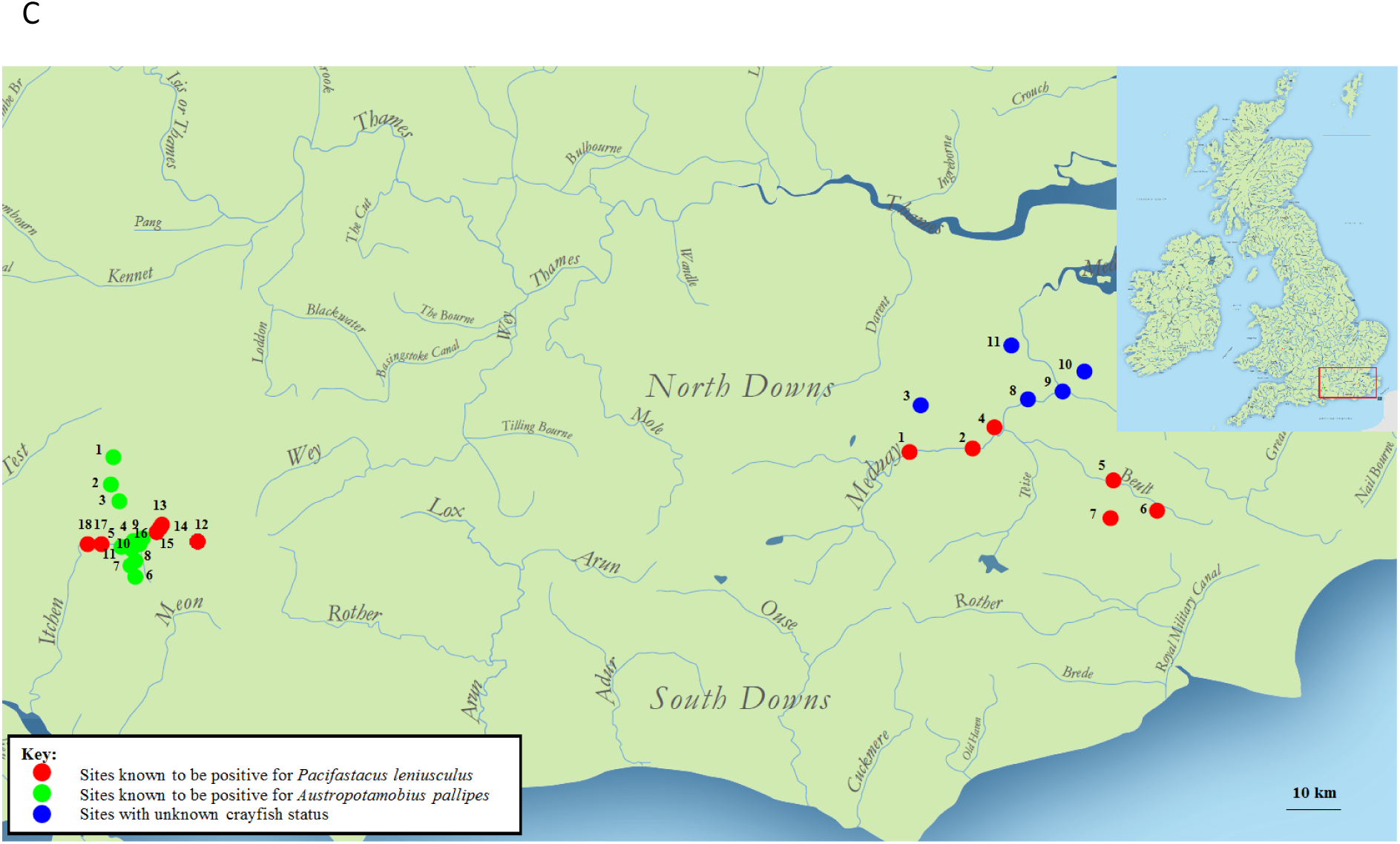
Locations of the rivers Itchen and Medway eDNA sampling sites. In the Itchen, there were 18 sites in total (I1 to I18); I1 – I11 classified as positive for *Austropotamobius pallipes* presence and I12 – 18 classified as positive for *Pacifastacus leniusculus* presence. In the Medway, there were 11 sites in total (M1 to M11); M1, M2 – M6 were classified as positive for *Pacifastacus leniusculus* presence whereas M3, M8 – M11 have an unknown crayfish species status. Each point corresponds to six water samples collected.

**Table 1.**
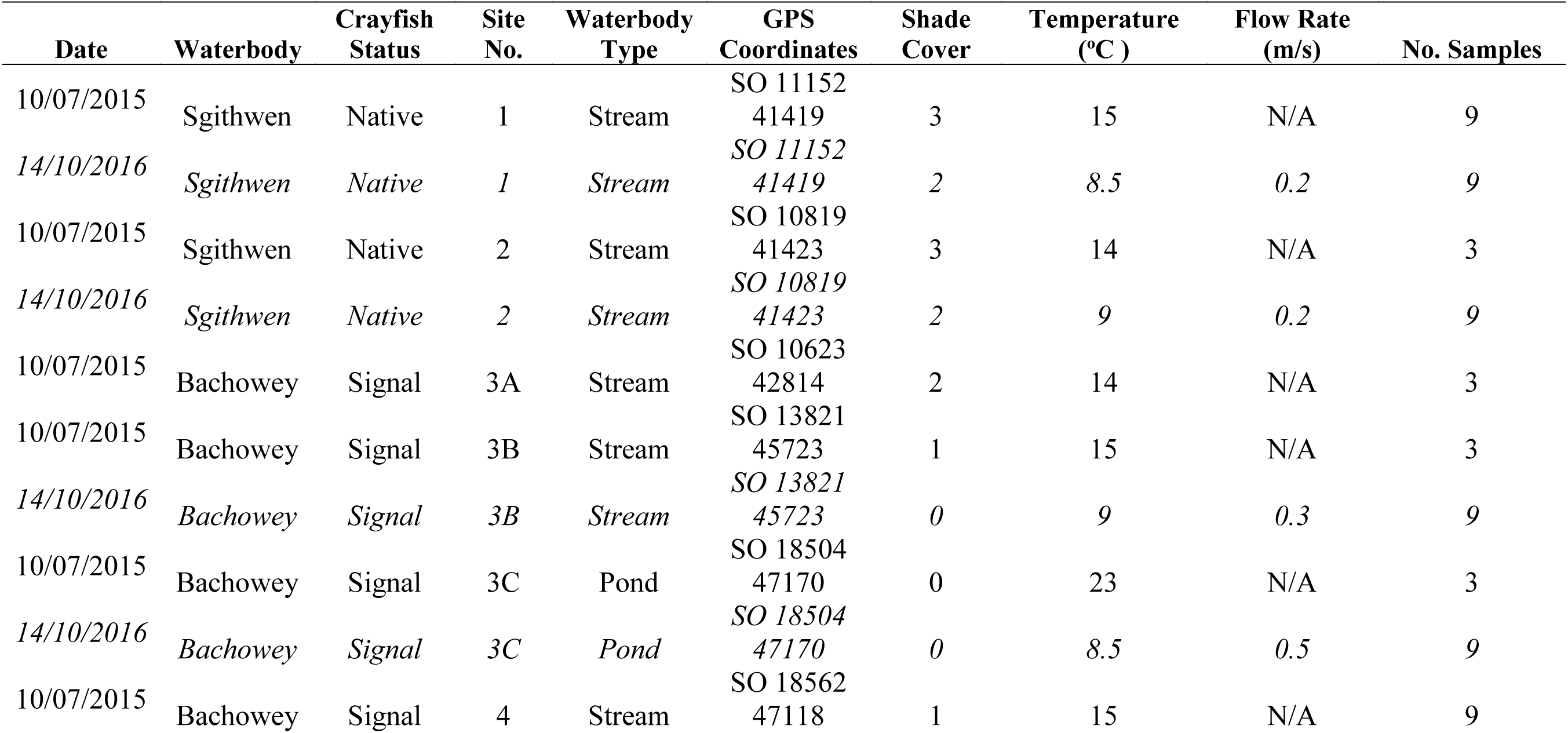
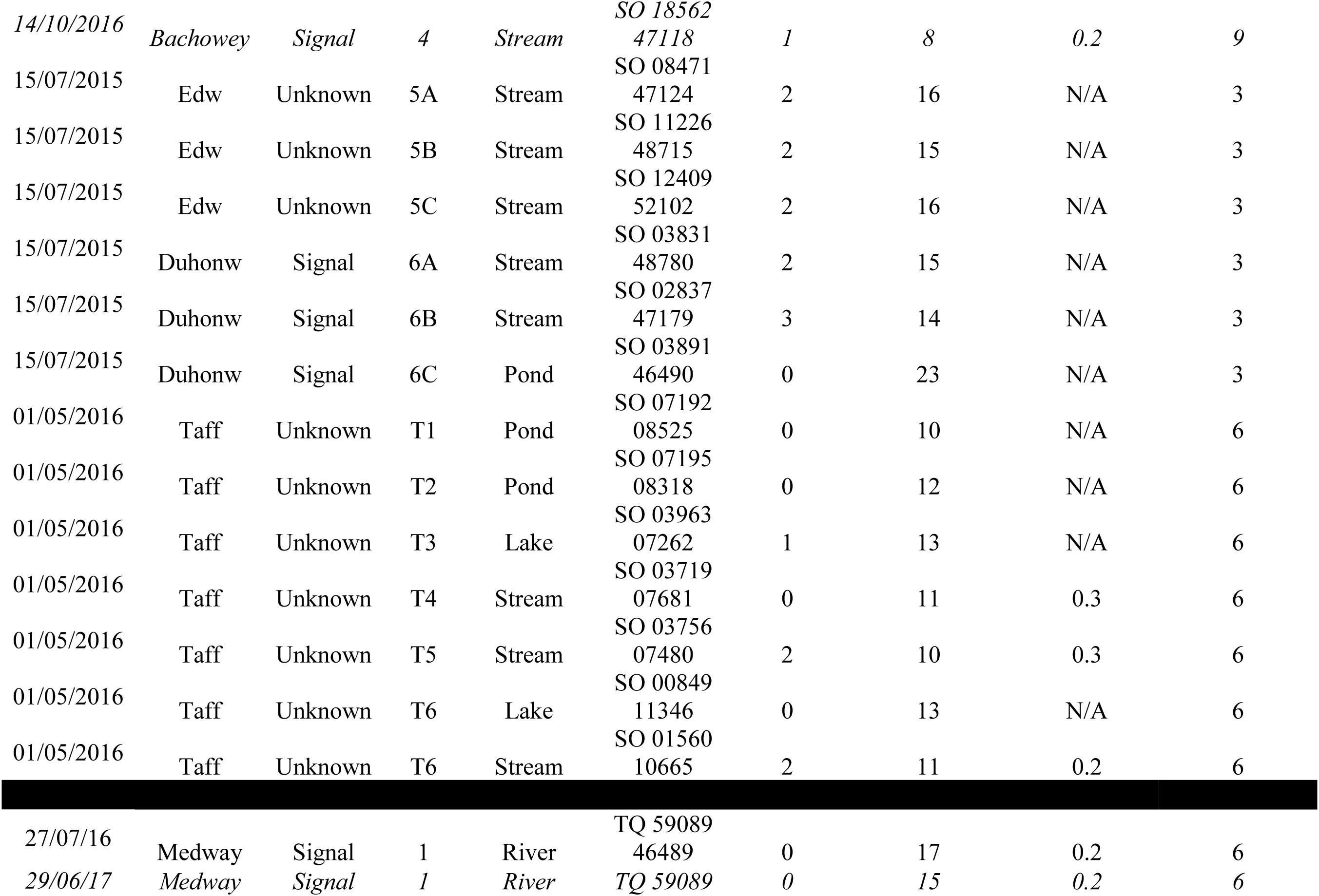
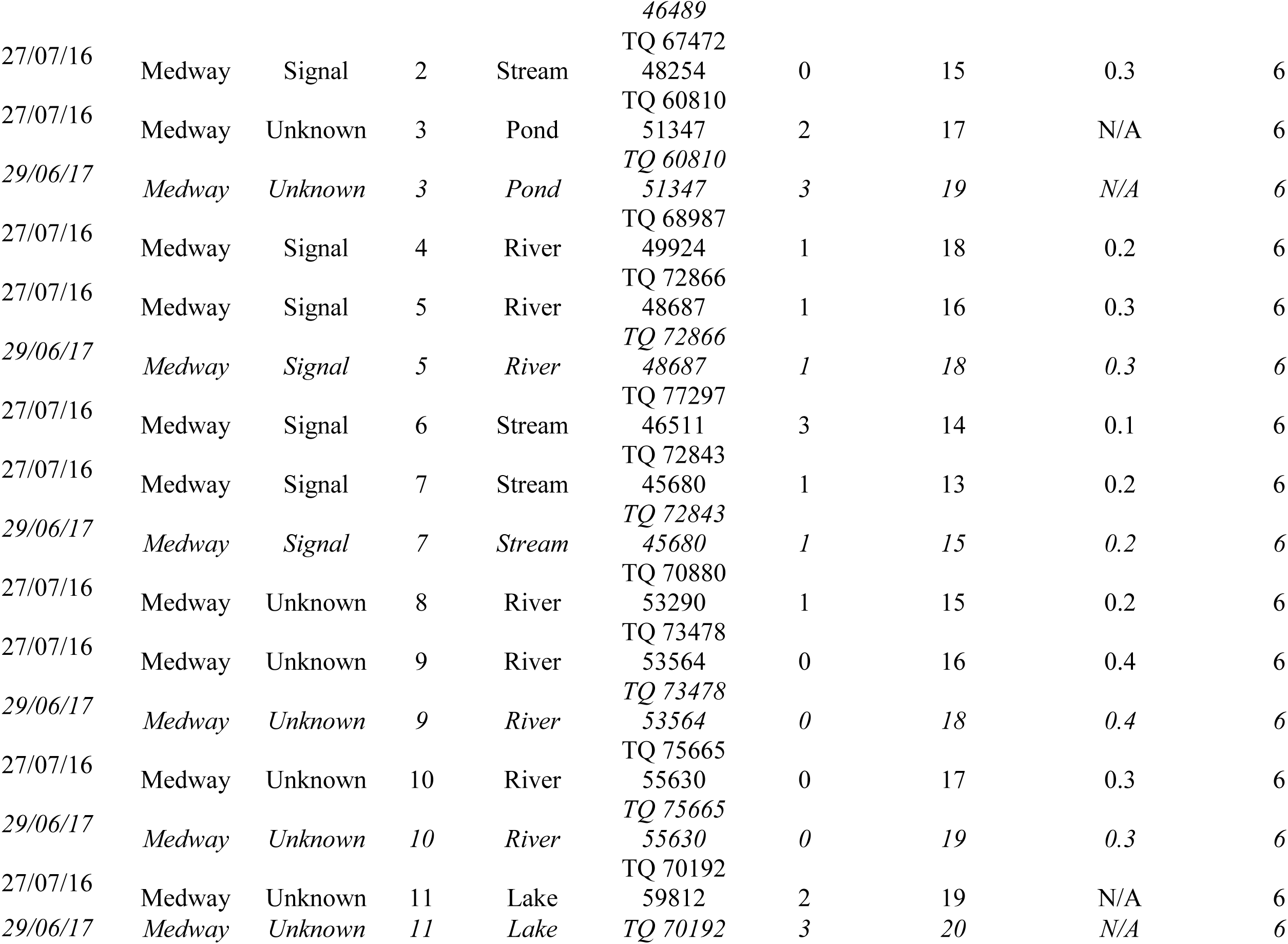
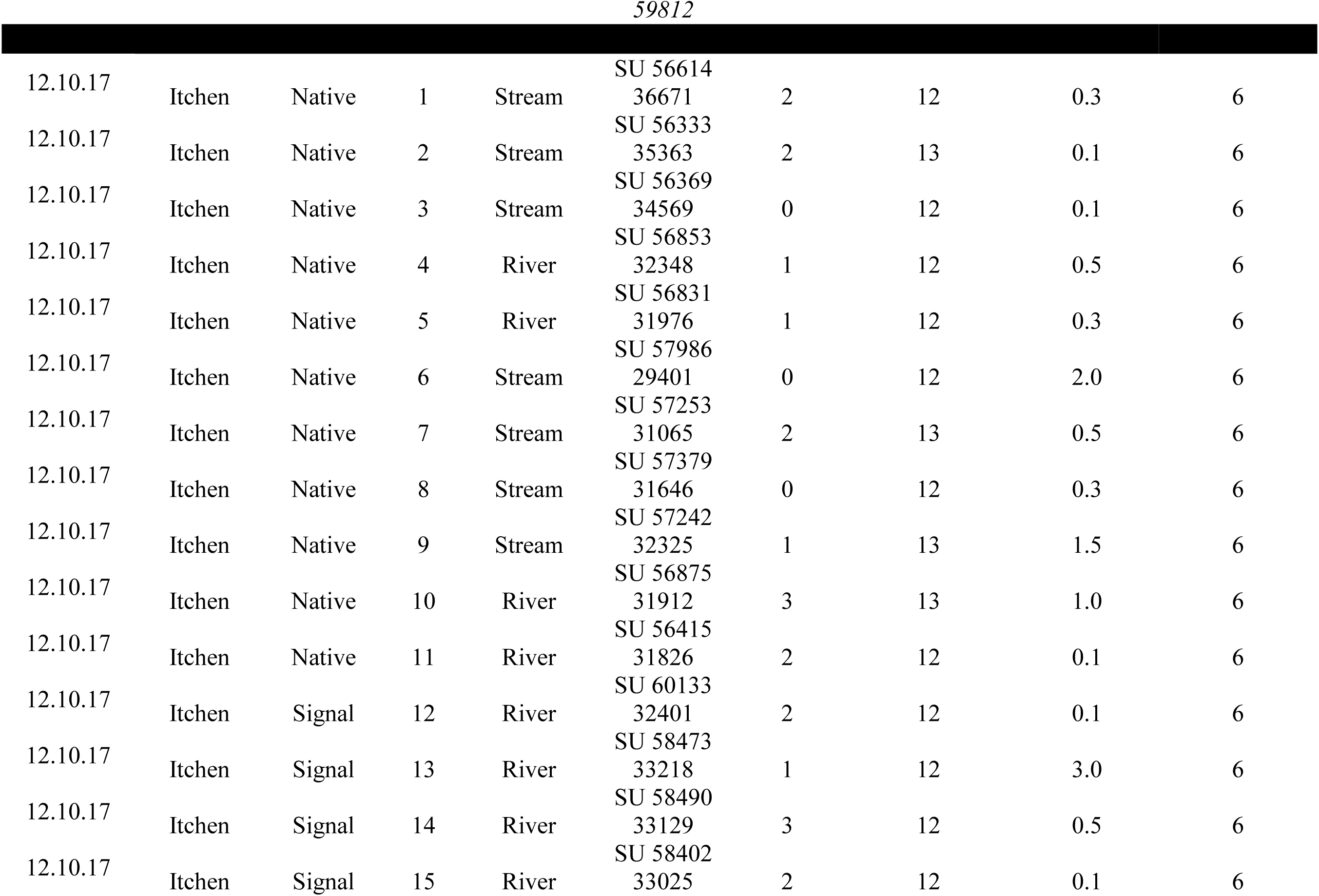
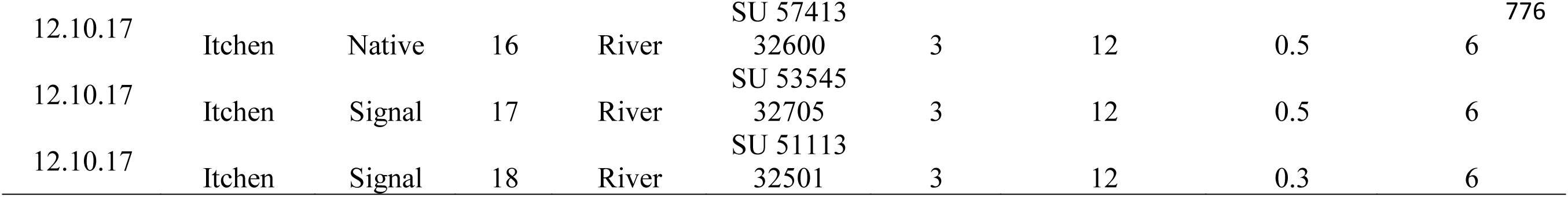
Location and environmental data for eDNA sampling sites in the River Wye for July 2015 and October 2016 (in italics); River Taff (May 2015); River Medway July 2016 and June 2017 (in italics) and the River Itchen (October 2017), including waterbody type, GPS coordinates, shade cover (0-3), temperature (°C), flow rate (m/s) and total number of samples collected per site minus negative controls (three samples in duplicate (6) or triplicate (9)).

Welsh locations were selected based upon data from CrayBase (James et al. 2014a); two of the locations supported *A. pallipes* populations, with no evidence of *P. leniusculus* presence, three locations only had populations of *P. leniusculus* and the remaining eight locations could potentially have both *P. leniusculus* and *A. pallipes* or neither species, but their status was uncertain as these had not been previously monitored. Two out of the three *P. leniusculus* confirmed sites were known to contain *A. astaci* infected crayfish (James et al. 2017).

In the river Medway, *P. leniusculus* was thought to inhabit the upper catchment but the crayfish status downstream was unknown, while in the river Itchen *A. pallipes* was assumed to be present throughout most of the upper catchment and *P. leniusculus* had been recorded in few sites both upstream and downstream of *A. pallipes* presence (Rushbrook 2014); Table 1). The infection status of both the Medway and Itchen crayfish populations was unknown.

Each site was subdivided into three sampling sites (upstream, midstream and downstream), separated where possible by ca. 500 m, to increase the area sampled. Between three and nine 15 mL water samples were taken from each sampling site simultaneously. All samples were collected ca. 1 m beneath the surface for ponds and in shallow areas of low flow streams and preserved as for the *ex-situ* experiment. Negative controls consisting of ultrapure water in place of river/pond water were taken before and after sampling, at each sampling site. Temperature, weather conditions, amount of shade cover, flow rate and pH were measured at each site (Table 1). Footwear was washed with Virkon™ and equipment disinfected with bleach between samplings to prevent the possible spread of *A. astaci* spores and DNA contamination between sites. All Wye sites which indicated presence of either crayfish species based on initial qPCR results were re-sampled the following year to assess reproducibility of positive amplifications at the sites (Table 1). To estimate the current presence of both host species, 25 standard TRAPPY™ crayfish traps (500 × 200 × 57 mm; NRW Permit Reference: NT/CW081-B-797/3888/02) were set following standard guidelines for trapping crayfish (DEFRA 2015). Traps were baited with halibut pellets and set at all of the eDNA sample sites and left for 24-48 hour, and 24 hour checks were conducted. Three 15 mL water samples were taken downstream of traps (or around the trap for still water bodies) which had successfully trapped crayfish, as a control of crayfish eDNA detectability in the river. Crayfish were collected and euthanised by freezing at -20 °C (Cooper 2011). Environmental data was recorded at each site as detailed above (Table 2).

**Table 2.**
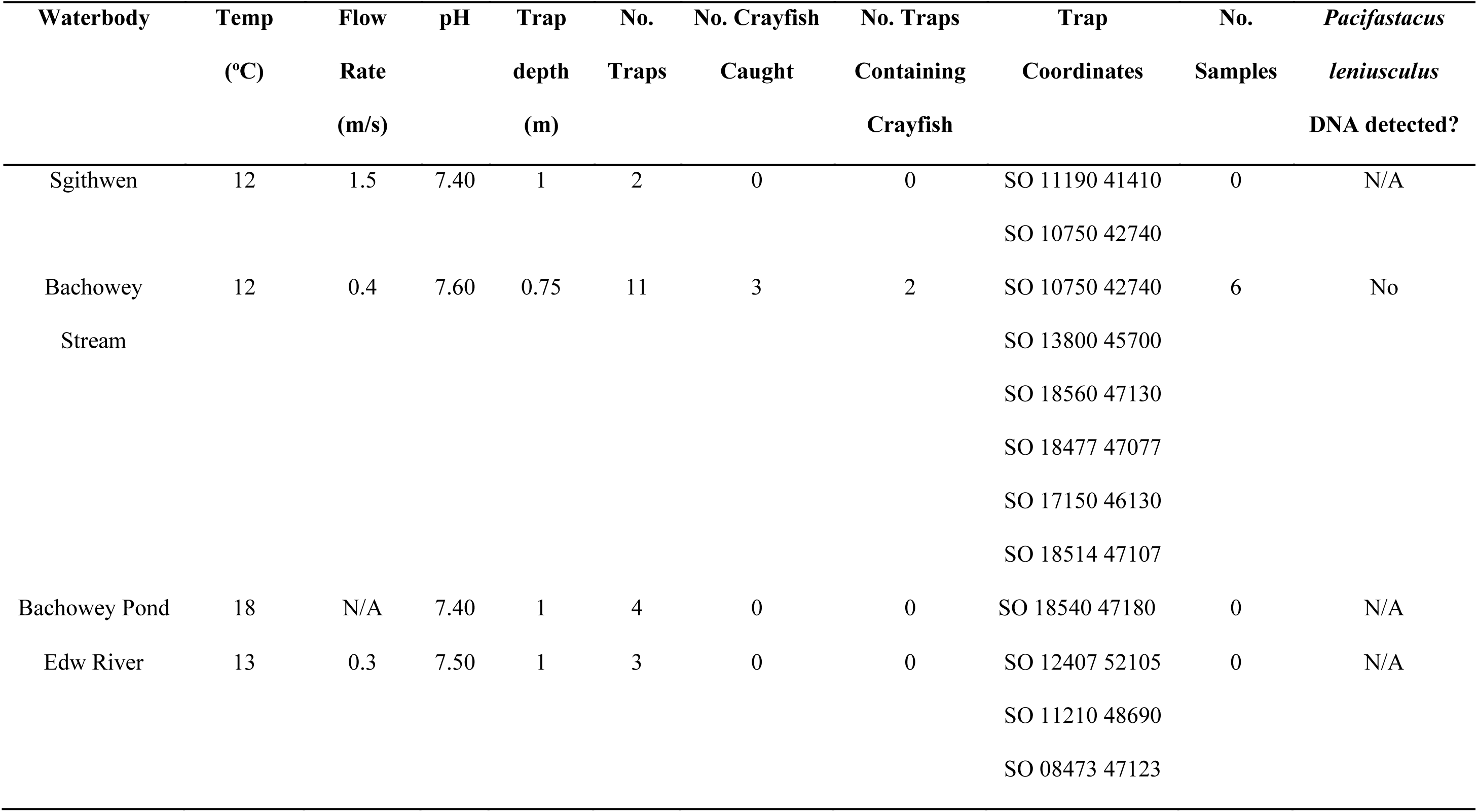
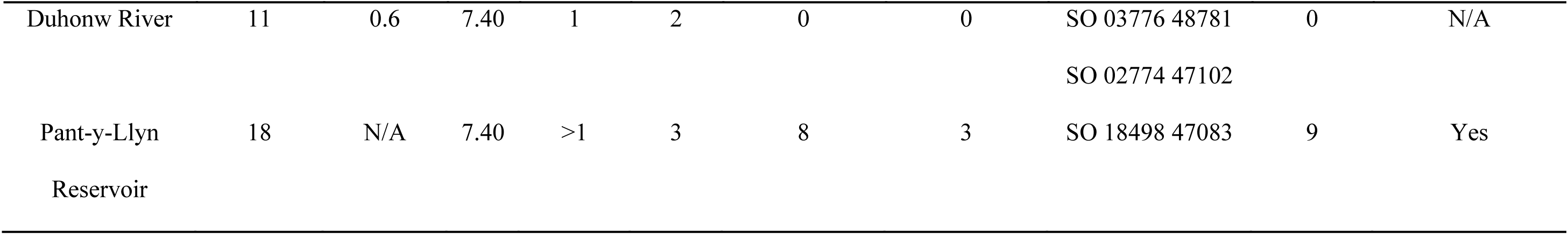
Location of crayfish traps in corresponding waterbodies in the Wye catchment and number of crayfish caught per trap.

Positive controls for eDNA screening consisted of 15 tissue samples from *P. leniusculus* individuals (pooled tail fan and soft cuticle) from three different source populations (Gavenny, Bachowey and Mochdre), part of a previous study within close proximity to eDNA sampling sites within the Bachowey and Duhonw catchments (James et al. 2017), and 12 *A. pallipes* individuals (first carapace moults and mortalities preserved in 100% ethanol) from two different locations in the UK (Cynrig Hatchery, Brecon and Bristol Zoo).

### qPCR PRIMER DESIGN

Crayfish specific primers were designed using Primer3 software, tested *in silico* using Beacon Primer Designer (ver. 2.1, PREMIER Biosoft), and checked for cross-amplification using NCBI Primer-BLAST (Ye et al. 2006). The primer pair was designed to be complementary to both the signal crayfish and native white-clawed crayfish (ApalPlen16SF: 5’-AGTTACTTTAGGGATAACAGCGT-3’ and ApalPlen16SR: 5’-CTTTTAATTCAACATCGAGGTCG-3’), to allow the amplification of an 83bp fragment of the 16S mtDNA gene (Data in brief Figure 1). The primers were assessed *in vitro* using positive control tissue (crayfish tail fan clips and moults) from 15 different signal and white-clawed crayfish individuals. DNA was extracted using Qiagen^®^ DNeasy Blood and Tissue Kit (Qiagen, UK), eluted in 100 µl, and amplified in end-point PCR using the following ApalPlen16S protocol: 95 °C for 3 min, followed by 40 cycles of 95 °C for 30 s, 61.5 °C for 30 s and 72 °C for 45 s with a final elongation step of 72 °C for 10 min. All amplified PCR products were checked for the correct amplicon size using a 2% agarose gel electrophoresis. Primers were also tested on tissue samples from a second invasive crayfish species established in the UK, the virile crayfish (*Orconectes* cf. *virilis*), and against a related species commonly found in the same environment, the freshwater shrimp (*Gammarus* sp.) to check for non-specific amplification.

DNA from the *ex-situ* eDNA samples for *P. leniusculus* and *A. pallipes* were extracted using Qiagen^®^ DNeasy Blood and Tissue Kit (Qiagen, UK), eluted in 100 µl, and amplified with ApalPlen16S primers. PCR products were run in a 2% agarose gel to check for correct amplicon size against positive controls (extracted crayfish tail clip), purified and analysed using Sanger Sequencing on an ABI Prism 277 DNA sequencer. Resulting sequences were aligned using BioEdit v. 5.0.9 (using the ClustalW program) and inputted to BLAST (Ye et al. 2006) to confirm the species identity.

### qPCR-HRM OPTIMISATION

Specific *in vitro* testing of RT-qPCR-HRM analysis was performed for both *P. leniusculus* and *A. pallipes* crayfish samples to ensure that each species could be identified based on their specific differential PCR product melt temperatures. Annealing temperature for ApalPlen16S primers was optimised at 61.5 °C and resulting efficiency values at this temperature for both species were 92.0 and 93.8% for *P. leniusculus* and *A. pallipes*, respectively. For optimisation, the ApalPlen16S-qPCR cycling protocol began with 15 min of denaturation at 95 °C, followed by 40 cycles of 95 °C for 10 s and 61.5 °C for 30 s. A HRM step was applied to the end of RT-qPCR reactions, ranging from 55 °C to 95 °C in 0.1 °C increments to assess the consistency of amplicon melt temperature (tm) for both crayfish species. Limit of detection (LOD) and limit of quantification (LOQ) were determined through running a dilution series ranging from 5 ng/µl to 5 × 10^−7^ ng/µl, using DNA pools for both species. HRM analysis was conducted on a minimum of 12 and a maximum of 15 individuals from several *P. leniusculus* and *A. pallipes* populations to account for any potential intraspecific variation in qPCR product tm (Table 3). qPCR-HRM analysis was undertaken comparing two master mixes, SYBR^®^ Green (Bio-Rad, UK) and SsoFast™ EvaGreen^®^ (Bio-Rad, UK), assessing consistency and reproducibility of both with relation to melt curve profiles (Table 3). To assess ability to detect both crayfish species in the same reaction, equal volumes of *P. leniusculus* and *A. pallipes* DNA were pooled together from ten different individuals of both species at various concentration ratios (ranging from 50:50 to 10:90).

**Table 3.**
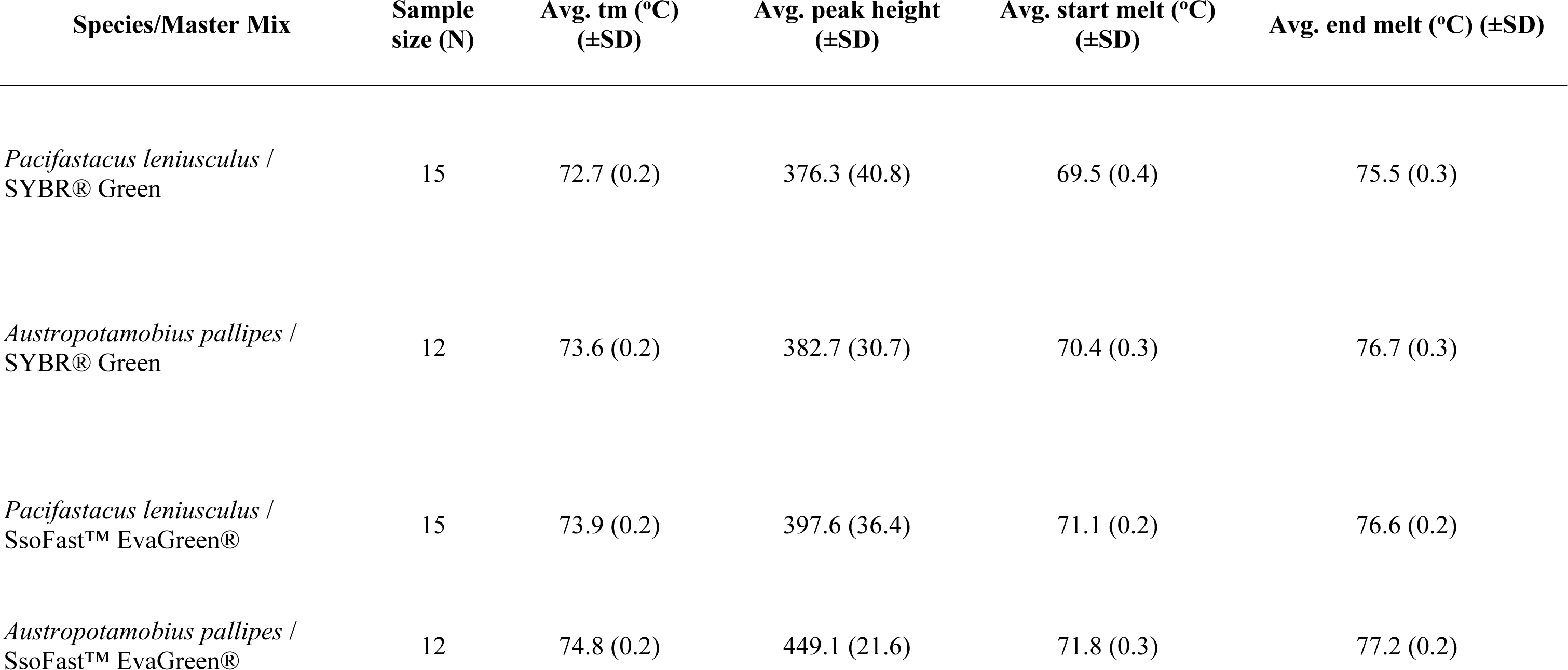
Summary of average values from qPCR outputs for both *Pacifastacus leniusculus* and *Austropotamobius* pallipes. Average melt temperature (°C; Avg. tm); Average melt peak height (Avg. peak height); Average start melt temperature (°C; Avg. start melt); Average end melt temperature (°C; Avg. end melt) of resultant qPCR products with standard deviation. Values were obtained for each individual over at least three separate runs, each consisting of three replicates and negative control blanks.

| Species/Master Mix | Sample size (N) | Avg. tm (°C) (±SD) | Avg. peak height (±SD) | Avg. start melt (°C) (±SD) | Avg. end melt (°C) (±SD) |
| --- | --- | --- | --- | --- | --- |
| *Pacifastacus leniusculus* / SYBR® Green | 15 | 72.7 (0.2) | 376.3 (40.8) | 69.5 (0.4) | 75.5 (0.3) |
| *Austropotamobius pallipes* / SYBR® Green | 12 | 73.6 (0.2) | 382.7 (30.7) | 70.4 (0.3) | 76.7 (0.3) |
| *Pacifastacus leniusculus* / SsoFast™ EvaGreen® | 15 | 73.9 (0.2) | 397.6 (36.4) | 71.1 (0.2) | 76.6 (0.2) |
| *Austropotamobius pallipes* /<br>SsoFast™ EvaGreen® | 12 | 74.8 (0.2) | 449.1 (21.6) | 71.8 (0.3) | 77.2 (0.2) |

Once the *in vitro* testing was complete for positive controls, further testing was undertaken for the eDNA samples collected in the *ex-situ* study to ensure that the primers would amplify environmental DNA samples and to assess the minimum levels of detection of eDNA samples.

### MULTIPLEX OPTIMISATION

For the *A. astaci* multiplex assay, optimisation of primer quantity and concentration was undertaken by combining the two sets of primers (ApalPlen16S and AphAstITS; (Vrålstad et al. 2009) at starting concentrations between 1 µM and 20 µM. Equal concentrations of each set of primers at 5 µM produced the most efficient co-amplification for both sets of primers, with poor amplifications resulting in concentrations from 1 to 4 µM and above 6 µM starting concentration. Uninfected crayfish DNA controls were obtained through extraction of a tail fan clip from non-infected individuals and *A. astaci-*positive samples were obtained from a previous study by Cardiff University (James et al. 2017), where an infected crayfish tail fan clip, melanised soft cuticle and walking leg tissue were pooled together and DNA extracted for *A. astaci* screening.

The final optimised multiplex qPCR reactions were carried out in a final volume of 10 µl, which contained 2 µl 5 x HOT FIREPol^®^ EvaGreen^®^ qPCR Mix Plus ROX (Soils Biodyne, Estonia), 0.4 µl of primer mix (5 µM), 1 µl template DNA at 5 ng/µl and 6.6 µl of ultrapure water. The amplification was carried out using a Bio-Rad CFX96 Touch Real-Time PCR Detection System (Bio-Rad, UK). The PCR protocol was as follows: once cycle of initial activation at 95 °C for 12 min, followed by 40 cycles of 95 °C for 15 s, 61.5 °C for 20 s and 72 °C for 20 s. After the PCR reaction, a melt curve program was set, which ran from 65 °C to 95 °C by raising 1 °C for 10 s each step. The resulting curve was then used to assess the presence/absence of *A. astaci* and target crayfish species DNA based on the species-specific melting temperatures of the DNA product (*A. astaci* = 82.9 °C; *P. leniusculus* = 75.9 ± 0.2ºC and *A. pallipes* = 76.6 ± 0.2ºC) which were identified during optimization of the multiplex assay.

### eDNA *IN SITU* ANALYSES

eDNA extraction from 407 field samples (Table 1) was performed using Qiagen^®^ DNeasy Blood and Tissue Kit (Qiagen, UK), following the manufacturer’s instructions, apart from a reduction in the elution volume from a single elution step of 200 µl to two elution steps of 50 µl to maximise DNA yield. DNA extractions took place in a dedicated eDNA area within an extraction cabinet, equipped with a UV light and a flow-through air system to minimise chances of contamination. Extractions were conducted wearing eDNA-dedicated laboratory coat, face mask and gloves. Samples were amplified in triplicate in a Bio-Rad CFX96 Touch Real-Time PCR Detection System (Bio-Rad, UK), in 10 µl reactions consisting of 5 µl SsoFast™ EvaGreen^®^ Supermix (Bio-Rad, UK), 0.25 µl each ApalPlen16SF and ApalPlen16SR, 3.5 µl HPLC water and 2 µl extracted DNA. Amplifications were carried out in triplicate using standard ApalPlen16s-qPCR protocol as described above and only samples which amplified consistently in at least two replicates at the target DNA product tm (either 73.9 ± 0.2 or 74.8 ± 0.2 °C), with a melt rate above 200 -d(RFU)/dT were considered to be a positive result. qPCR reactions were carried out in a separate room to eDNA extractions under a PCR hood with laminar flow. Two positive controls per species were added to each plate once all the eDNA samples were loaded and sealed to prevent false positives in the eDNA samples. Two amplification negative controls consisting of HPLC water and two extraction negative controls were also added in the same well location on each plate test for contamination in eDNA samples.

A subset of positive field samples, along with a positive control for each crayfish species were re-amplified using end-point PCR, purified and cloned into pDrive plasmid cloning vector (Qiagen PCR Plus Cloning Kit, Qiagen, UK). Three to nine clones per sample were sequenced using T7 and SP6 primers on an ABI Prism 377 sequencer.

## RESULTS

### *EX-SITU* OPTIMISATION

Optimisation of eDNA protocols was carried out *ex-situ* by placing individual *P. leniusculus* in isolated tanks for 24 hours and sampling water from those tanks 24 and 48 hours after removal. Reference DNA from the *ex-situ* study was successfully extracted and amplified in triplicate from *P. leniusculus* and *A. pallipes* positive controls and species confirmed by Sanger Sequencing of the 83bp fragment of the 16S mtDNA. DNA from signal crayfish was detected in all water samples taken at different time points from the *ex-situ* study. eDNA concentrations marginally decreased overtime and correlated with Cq values for *ex-situ* samples in qPCR amplifications (Data in brief: Figure 2; Figure 5B; Table 3). DNA from native crayfish was also amplified in all nine water samples from the reference hatchery. No amplification bands were present in any of the negative controls (tank, extraction and amplification).

**Figure 2.**
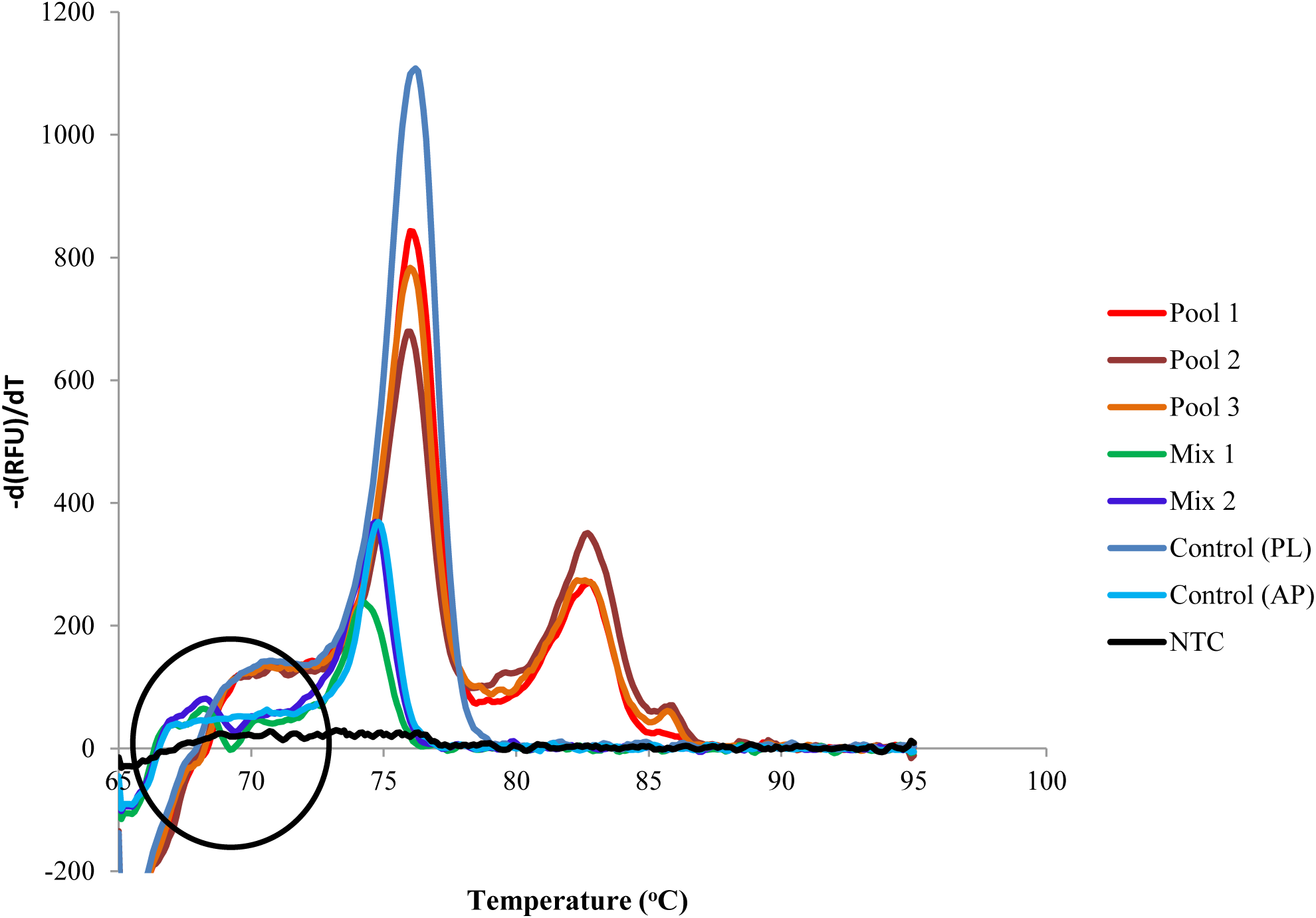
qPCR product melt peak output for multiplex amplification of DNA using optimised HOT FIREPol^®^ EvaGreen^®^ from three different *Pacifastacus leniusculus* individuals and *Aphanomyces astaci* DNA in the same qPCR reaction (Pool 1-3), displaying the diagnostic double melt peaks at 75.9 ± 0.2 °C for *Pacifastacus leniusculus* and 82.9 °C for *Aphanomyces astaci*. Also displayed, is the qPCR melt profiles for mixed proportions of *Pacifastacus leniusculus* and *Austropotamobius pallipes* DNA in the same qPCR reaction, showing the diagnostic melt shape difference between the three types of amplification (circled) and positive controls. Mix 1 = 90:10 *Pacifastacus leniusculus*: *Austropotamobius pallipes*, Mix 2 = 50:50 *Pacifastacus leniusculus*: *Austropotamobius pallipes*, PL = *Pacifastacus leniusculus* and AP = *Austropotamobius pallipes*.

### CRAYFISH DETECTION LIMITS

The results of the qPCR optimisation indicated that the limit of detection (LOD) of both *P. leniusculus* and *A. pallipes* DNA was 0.005 ng/µl, after a 10-fold dilution series. The detection threshold for amplification of positive control DNA used for optimisation from both species was between 16 and 28 cycles, and the melting temperatures (tm) of the DNA products were consistent for both *P. leniusculus* and *A. pallipes*, with no overlap between the two species (Table 3). SsoFast™ EvaGreen^®^ multiplex master mix performed more consistently than the SYBR^®^ Green master mix, with a lower standard deviation for average tm, average peak height, average start melt temperature and average end melt temperature (Table 3; Data in brief: Figure 3; Table 1). Results of the qPCR analysis of mixed proportions of *P. leniusculus* and *A. pallipes* DNA confirmed that it is possible to discriminate between positive amplifications of eDNA for single crayfish species vs. mixed crayfish species (*P. leniusculus* and *A. pallipes*). Diagnostic peaks in early product melt temperatures were present for all amplifications containing 90:10 to 50:50 ratios of *P. leniusculus: A. pallipes* DNA (Figure 2; Data in brief: Figure 5A; Table 2).

**Figure 3.**
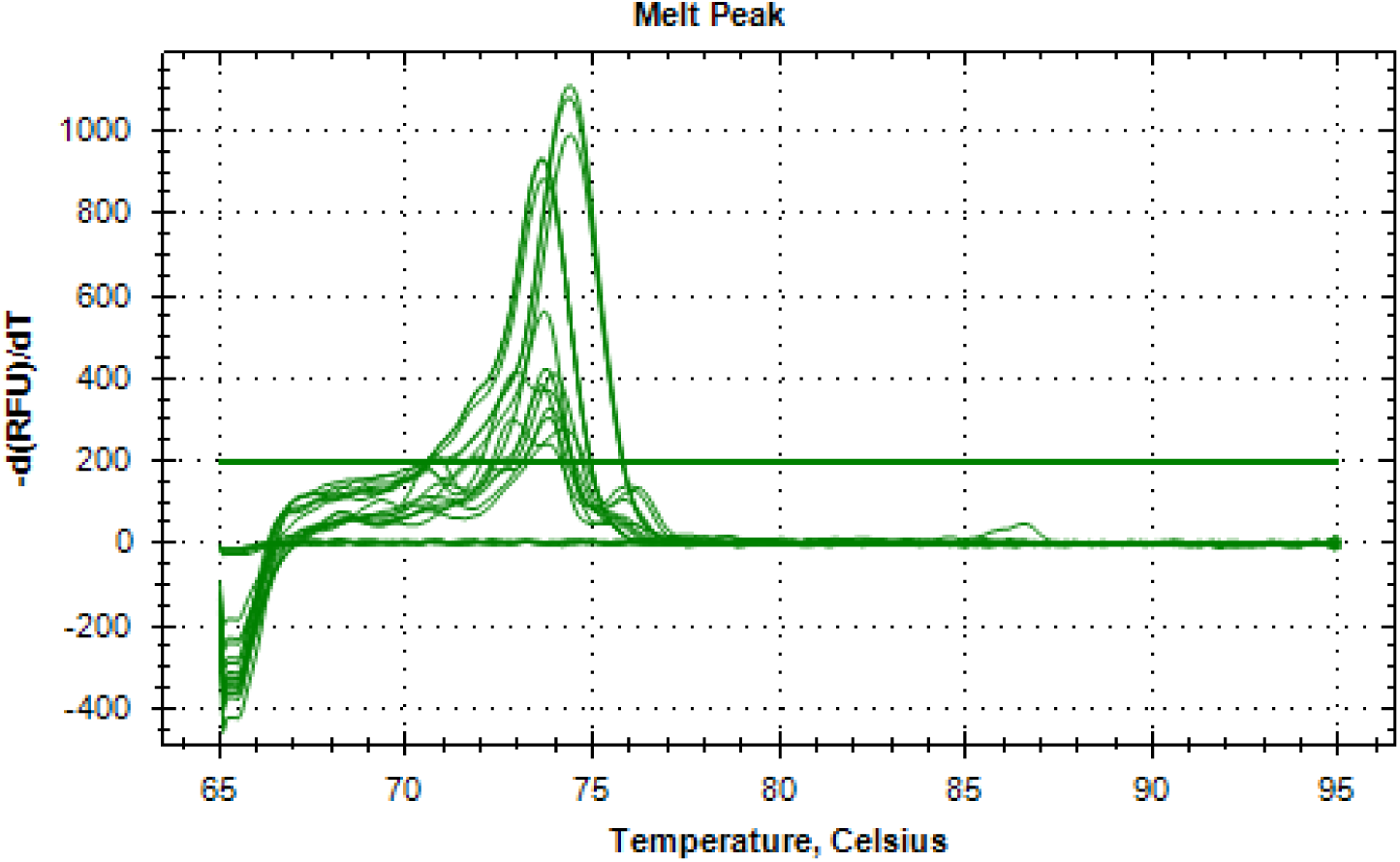
Melt peak profile for HOT FIREPol^®^ EvaGreen^®^ multiplex amplification of samples 5B, 5C and 5D from site 5 in the Taff catchment. The two largest sets of peaks correspond to positive control tissue for both *Pacifastacus leniusculus* (73.7 °C) and *Austropotamobius pallipes* (74.7 °C) and subsequent peaks represent eDNA field sample melt peaks for both *Pacifastacus leniusculus* and *Austropotamobius pallipes*. Non-template had no melt profile (flat line).

### SIMULTANEOUS DETECTION OF CRAYFISH AND *APHANOMYCES ASTACI*

The multiplex assay for simultaneous crayfish and *A. astaci* detection resulted in two products with an average tm of 75.9 ± 0.2 ºC for *P. leniusculus* (or 76.6 ± 0.2 ºC for *A. pallipes*; four individuals) and 82.9 °C for *A. astaci.* DNA controls from four *A. astaci*-infected *P. leniusculus* individuals (INF 1 – INF 4) were successfully amplified with two products of the corresponding temperatures. Amplification of uninfected *P. leniusculus* DNA resulted in a single product with tm of 75.9 ± 0.2 ºC (Figure 2; Data in brief: Figure 6; Table 4).

**Table 4.**
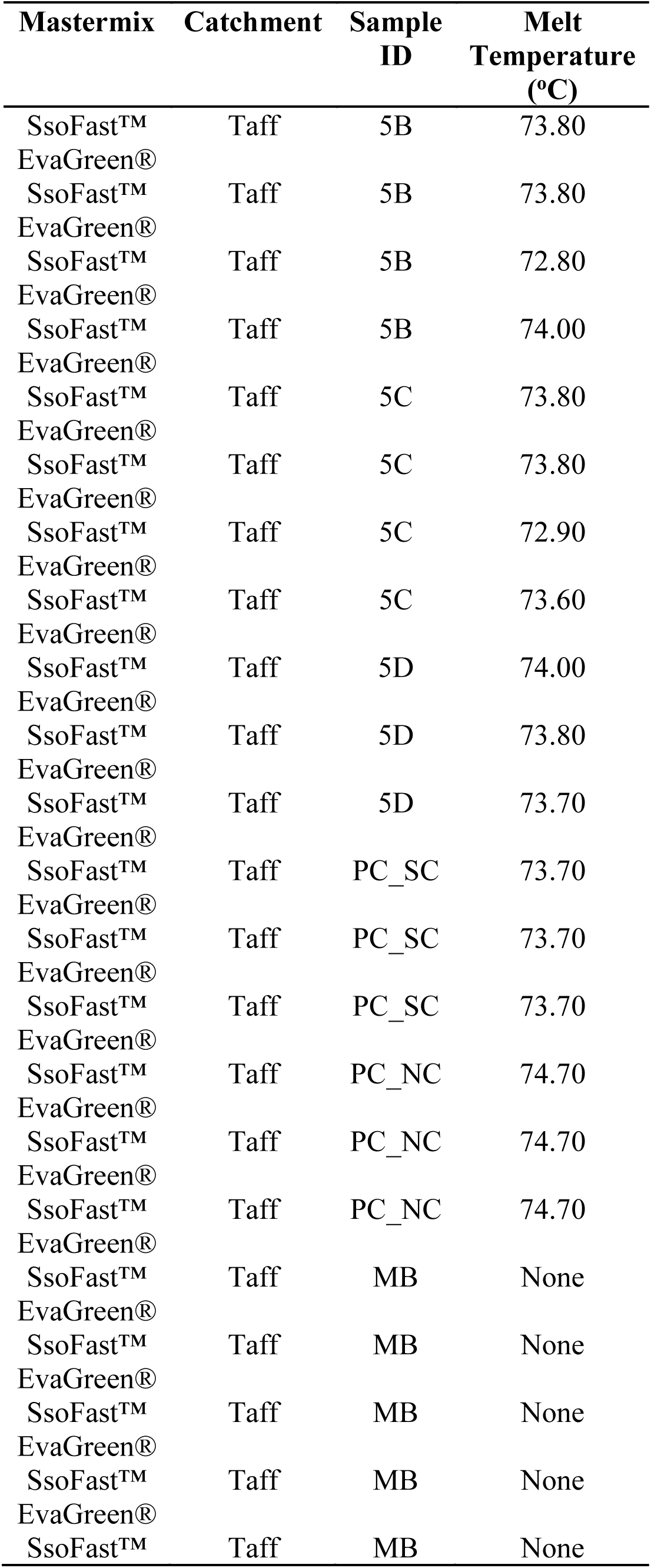
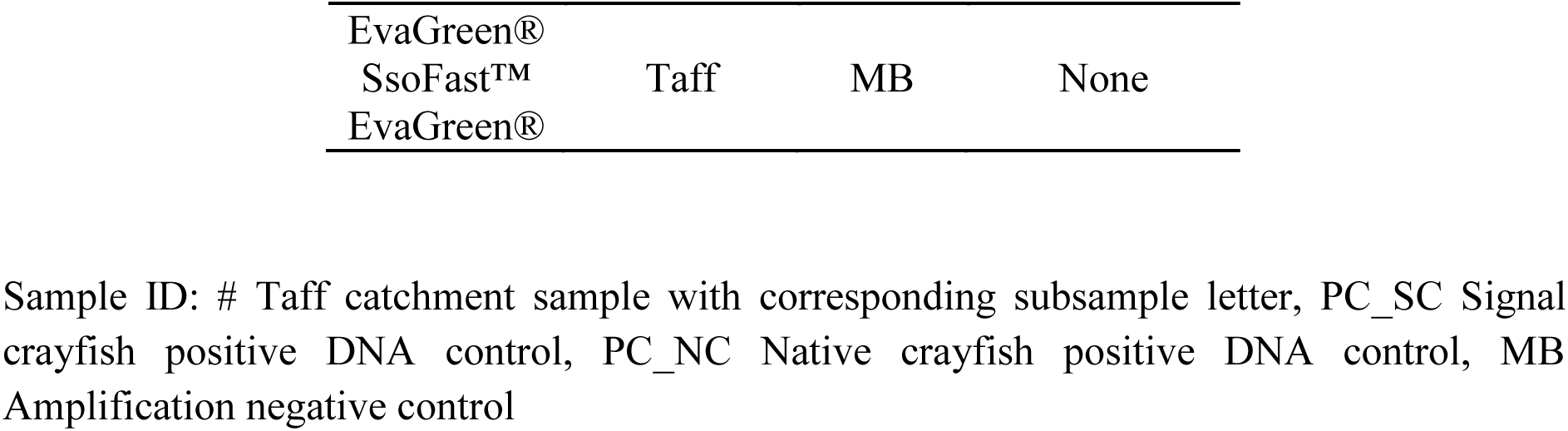
Melt data from SsoFast™ EvaGreen^®^ eDNA qPCR amplifications for the Taff catchment.

### CRAYFISH SPECIES DISTRIBUTION AND INFECTION STATUS

For Welsh sites, crayfish trapping confirmed the presence of *P. leniusculus* (11 caught across 3 different sites; Table 2) in positive sites, whereas no *A. pallipes* were caught, despite visual confirmation of the species upon collecting traps. *P. leniusculus* eDNA was successfully detected around each of the three traps in the reservoir (Data in brief: Figure 7; Table 5). qPCR detected *P. leniusculus* eDNA at all three confirmed sites for the species and *A. pallipes* eDNA was detected within the confirmed tributary for the species (Data in brief: Figure 8A-B; Table 6). Additionally, *P. leniusculus* eDNA was detected in one of the unknown crayfish status sites in the river Taff whereas there was no positive detection of *A. pallipes* in any of the other the sites with unknown presence of the species (Figure 3, Table 4).

In both the Medway and Itchen there was evidence of *P. leniusculus* and *A. pallipes* coexisting in two sampling sites (Data in brief: Figure 9; Table 7). One site in the Medway was positive for both crayfish species over the two-year sampling period and one site in the Itchen was also positive for both species in the single sampling event carried out. Both *A. pallipes* and *P. leniusculus* were also detected in the Medway and Itchen in separate areas (*A. pallipes*: Medway (2 sites), Itchen (4 sites); *P. leniusculus*: Medway (3 sites), Itchen (9 sites).

A. *astaci* was confirmed in all sites in the river Bachowey, resulting in two products with melt peaks at 75.9 ± 0.2 and 82.9ºC for the signal crayfish and plague agent respectively (Data in brief: Figure 8C; Table 6). All other sites positive for *P. leniusculus* or *A. pallipes* were negative for *A. astaci,* which was not detected in the rivers Medway or Itchen, despite the coexistence of both crayfish species (Figure 4).

**Figure 4.**
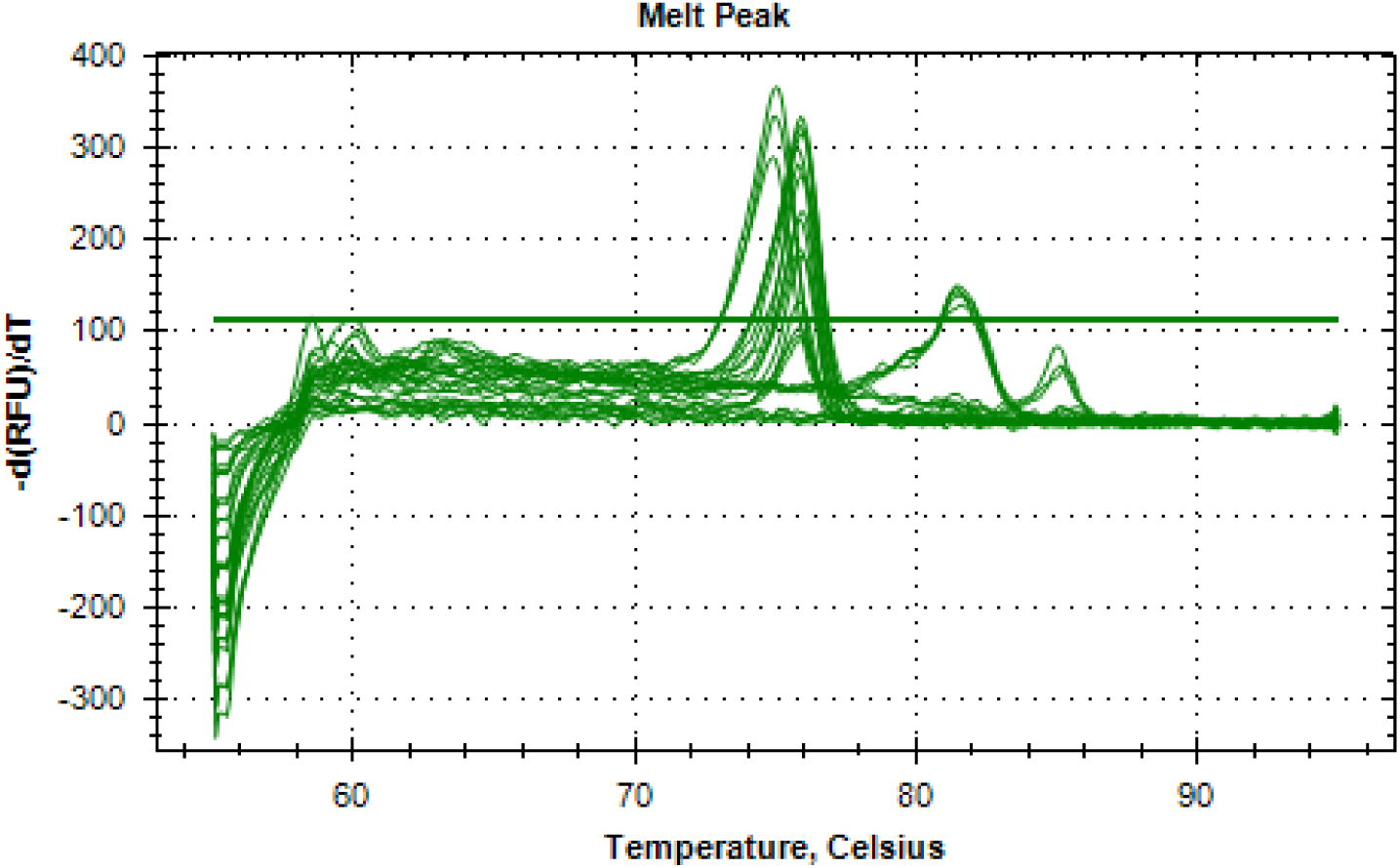
Melt peak profile for SsoFast™ EvaGreen^®^ eDNA qPCR amplifications of positive amplifications for both *Pacifastacus leniusculus* and *Austropotamobius pallipes* in the same site. The three largest sets of peaks correspond to positive control tissue (one sample in triplicate) for both *Pacifastacus leniusculus* (74.9 °C), *Austropotamobius pallipes* (75.9 °C) and *Aphanomyces astaci* (82.9 °C). Subsequent peaks represent eDNA field sample melt peaks from nine samples (in triplicate) for both native *Austropotamobius pallipes* and invasive *Pacifastacus leniusculus*, with absence of any melt peak for *Aphanomyces astaci* in field samples. Non-template control has no melt profile (flat line).

A subset of five positive amplifications was selected (one for *A. pallipes* and four for *P. leniusculus*) to confirm species identity by cloning and sequencing. Out of 36 successfully transformed clones for the field samples of *P. leniusculus* (nine for each sample), between two and nine clone sequences per sample matched 100% with *P. leniusculus* on BLAST (Ye et al. 2006); remaining clones were a product of non-specific amplification. For *A. pallipes* field samples, two out of 3 clones from the positive field sample matched 100% for *A. pallipes*. All six positive control clones matched 100% with respective crayfish species (*P. leniusculus*/*A. pallipes*).

## DISCUSSION

By using a novel multiplex approach we could simultaneously detect the presence of the endangered white clawed crayfish and the highly invasive North American signal crayfish within a catchment that was free of crayfish plague. In contrast, we did not detect any native crayfish or coexistence of both species in tributaries where the pathogen was identified. A common impact of invasive species on native populations is the transmission of pathogens. Many non-native species not only introduce novel pathogens (Miaud et al. 2016) but also act as non-clinical carriers, facilitating their dispersal (Andreou et al. 2012). In this way, pathogens can act as biological weapons that allow invasive species to outcompete their native counterparts (Vilcinskas 2015), as in the case of the UK native crayfish, highly susceptible to the plague carried out, mostly asymptomatically, by the invasive signal crayfish (Andreou et al. 2012). As highlighted in the principles adopted by the Convention on Biological Diversity on invasive species, prevention and early detection should represent the priority responses to invasive species to allow for rapid response and more cost-effective removal when possible (Simberloff et al. 2013) and our study is the first one to combine eDNA and HRM for early detection of novel pathogens carried by non-native species, being particularly relevant for management and conservation in relation to aquatic biological invasions.

The causal agent of crayfish plague, *A. astaci*, is considered one of the primary causes for the extirpation of native crayfish populations across Europe (Alderman et al. 1990; Dunn et al. 2009). Attempts to eradicate established populations of *P. leniusculus* and other invasive non-native crayfish have been largely unsuccessful and costly (Dougherty et al. 2016; Kirjavainen and Sipponen 2004; Peay 2009; Sandodden and Johnsen 2010) and increasing emphasis is being placed on early detection of non-native crayfish, rather than on eradication of established populations (Freeman et al. 2010; Gherardi et al. 2011; James et al. 2014b; Tréguier et al. 2014). Our protocols followed the most updated guidelines for the use of eDNA for aquatic monitoring (Goldberg et al. 2016), ensuring the consistency of our results. We first validated our method with positive controls and by detecting both native and signal crayfish in sites where they had been previously observed as well as detecting *A. astaci* in a recognised infected river.

Only native or invasive crayfish (not both species coexisting) were expected in the Wye catchment, where some populations of *P. leniusculus* are known to be carriers of the plague and have been established for a sufficient amount of time to entirely displace native *A. pallipes* from most of the species’ historical locations (Dunn et al. 2009; James et al. 2014b), and this was supported by our results. Our multiplex approach successfully identified *A. astaci* in the Bachowey stream and *P. leniusculus* in an associated pond less than 10 m from this stream, revealing the presence of infected crayfish further upstream than previously detected (James et al. 2017), despite previous intensive trapping of *P. leniusculus*, which removed 36,000 individuals from the area between 2006 and 2008 (Wye & Usk Foundation 2012). We also detected the endangered crayfish *A. pallipes* in spite of its very low abundance in the Sgithwen, made apparent by lack of trapping success, highlighting the sensitivity of the method.

In the rivers Medway and Itchen, where invasions date back to the 1970s (NBN 2009), both *P. leniusculus* and *A. pallipes* had been previously reported but the crayfish plague status was unknown. We did not find *A. astaci* DNA in any samples from either catchment but found both the native and the invasive species coexisting in at least two sampling sites. This could be explained by the absence of plague, as *A. astaci* is often the main cause of *A. pallipes* population declines (Haddaway et al. 2012). We consistently detected both species over two sampling periods in the Medway, highlighting the reproducibility of the results, which combined with the absence of crayfish plague DNA presence suggests this could be a location where both species’ populations are stable (Bubb et al. 2005; Kozubíková et al. 2008). Populations of *A. pallipes* and *P. leniusculus* can coexist for a substantial length of time (c.25 years), as has been observed in other invasive-native crayfish population assemblages (Kozubíková et al. 2008; Peters and Lodge 2013; Schrimpf et al. 2012), providing that there is no introduction of *A. astaci* (Kozubíková et al. 2008; Schrimpf et al. 2012). However, due to competitive exclusion, it is unlikely that populations of both species will coexist indefinitely (Schrimpf et al. 2012; Westman et al. 2002), therefore areas where they overlap should be prioritised for management and control of the invasive species.

Detectability was variable among sampling seasons. There were more positive *P. leniusculus* field samples from the sampling of Wye sites in October 2016 compared to the samples collected in July 2015 from the same sites, with three and one positive samples respectively. For *A. pallipes*, the only positive field samples for the Welsh sites were from samples collected in October, however eDNA from both *P. leniusculus* and *A. pallipes* was successfully detected in the Medway samples collected in June and July. Seasonal differences could be due to the influence of temperature on eDNA detection rates among aquatic species; with every 1.02 °C rise in temperature, species are 1.7 times less likely to be detected, especially if the populations are at very low abundance (Moyer et al. 2014), whereas time since DNA release seems to have less effect on detectability at constant temperature (Eichmiller et al. 2016; Moyer et al. 2014). As temperatures in the Wye catchment were around six degrees colder in the stream sites and up to 14 degrees colder in still water bodies in October compared to July, this could explain the differences in detection success among samplings in the Wye catchment (Eichmiller et al. 2016; Moyer et al. 2014). However, temperatures in the Medway were similar to those in the Wye in July suggesting that the differences in detectability between catchments could be due to differences in population size or to local environmental conditions increasing DNA degradation rates in the Wye (Barnes et al. 2014; Dougherty et al. 2016; Jane et al. 2015; Pilliod et al. 2014). In contrast, *A. astaci* sporulation occurs most efficiently at temperatures nearer 20 °C, which could result in more spores being present in the river system in the summer months in comparison to any other time of the year (Wittwer et al. 2018). Released zoospores can only survive up to three days without a host and encysted spores survive up to two weeks in water, particularly during summer months when average temperatures of flowing and enclosed waterbodies are above 18 °C (Diéguez-Uribeondo et al. 1995; Unestam 1966), meaning it is possible to achieve a relatively real-time picture of *A. astaci* prevalence in eDNA samples (Wittwer et al. 2018). Lower abundance of *A. astaci* spores in colder temperatures could explain lack of detection of *A. astaci* in the October samples at the positive July sites in the Wye catchment (Strand et al. 2014; Wittwer et al. 2018), although detection levels could also have been affected by natural variation in population levels of plague infection (James et al. 2017). Considering this variability, seasonal samplings repeated over at least two years are advisable to reliably map the presence/absence of native and invasive crayfish and determine their infectious status.

In contrast to other essays developed for crayfish detection (Agersnap et al. 2017; Cai et al. 2017; Dougherty et al. 2016; Mauvisseau et al. 2018), our single, closed tube reaction, reduces not only the processing time and number of reactions but also the risk of contamination inherent to carry out a larger number of amplifications. HRM has already proved highly specific and useful for multiple species identification (Naue et al. 2014) and for the management of terrestrial invasive species (Ramón-Laca et al. 2014) but had never been applied to the detection of aquatic invasive species and their impacts using eDNA. Implementation of our multiplex assay provided three-fold biological information (invasive/native/pathogen) on target species’, which allows to assess potential contributing factors to native crayfish decline (such as the presence of invasive crayfish and crayfish plague) with greater sensitivity, specificity and efficiency than trapping (Barnes and Turner 2015) or single-species assays, essential to inform effective conservation and management strategies (Darling and Mahon 2011; Kelly et al. 2014).

While most studies have mainly focussed on crayfish eDNA detection in closed systems (Agersnap et al. 2017; Cai et al. 2017; Dougherty et al. 2016; Mauvisseau et al. 2018), our method has also proved useful for monitoring in flowing water bodies. This is important for early detection of invasive crayfish which use rivers and streams as a means for dispersal (Bubb, Thom and Lucas 2004), and particularly for *A. pallipes* whose detection was marginally better using eDNA (7%) than trapping (0%). In terms of sampling effort, eDNA tends to be more time effective than trapping (Smart et al. 2015). However, we failed to detect crayfish in the deep reservoir at Pant-Y-Llyn using eDNA, where trapping had revealed the presence of *P. leniusculus*. Taxonomic groups such as fish and amphibians shed significantly more DNA into the environment compared to invertebrate species, especially those with a hardened exoskeleton such as the crayfish (Thomsen et al. 2012; Tréguier et al. 2014). This reduced release of extracellular DNA can lower the detectability of crayfish, resulting in an increased occurrence of false negatives (Ikeda et al. 2016), particularly when the concentration of DNA is low due to few individuals or large water volumes (Tréguier et al. 2014). The nature of the crayfish exoskeleton combined with the depth of the reservoir prevented samples being taken near the sediment where the crayfish reside could account for observed lack of detection at the Pant-y-Llyn site (Tréguier et al. 2014). Collection of sediment samples in addition to water could improve levels of detection of target species, because DNA from sediment can last longer and be more concentrated than in water (Turner et al. 2015).

Conservation efforts rely on efficient, standardised methods for collecting biological data, which advance beyond the limitations of traditional sampling methods (Thomsen and Willerslev, 2015). Ecosystem management and conservation strategies strive to protect biodiversity through preventing invasions from novel species (thus the need for early detection) and effectively monitoring rare native species to preserve hotspots and ark sites (Lodge et al. 2012). Environmental DNA has been directly used as a conservation tool to survey both invasive (e.g.Takahara et al. 2013; Tréguier et al. 2014) and endangered native species (e.g. Olson et al. 2012; Sigsgaard et al. 2015) and we have shown how an eDNA-based qPCR-HRM multiplex approach can identify invasive hosts and their pathogens as well as refugia for the native species. This was particularly important to identify areas of coexistence between aquatic native and invasive crayfish (e.g. at the early stages of invasion or where crayfish plague is absent) (Schrimpf et al. 2012), which could be prioritised for long-term conservation plans.

Incorporating this tool to monitoring programmes for conservation significantly reduces the costs of sample processing compared to species’ targeted methods. Our method can ultimately help in the early detection and prevention of dispersal of invasive hosts and pathogens in threatened freshwater ecosystems, as well as in determining suitable locations for the potential reintroduction of the native species to historic habitats. As genomic technology advances, environmental DNA assays should continue to provide additional information, including more accurate data on species abundance and biomass in both lotic and lentic systems (Bohmann et al. 2014; Rees et al. 2014) as well as development of additional multiplexes to simultaneously detect numerous target species of conservation interest.

## ACKNOWLEDGEMENTS

This research was funded by the Welsh Government and Higher Education Funding Council for Wales (HEFCW) through the Sêr Cymru National Research Network for Low Carbon Energy and Environment (AQUAWALES; NRN-LCEE) and by the Environment Agency UK. We thank: Jennifer Nightingale, Oliver Brown and Adam Petrusek for crayfish/DNA samples, Stephen Marsh-Smith, Louis MacDonald-Ames & Hayden Probert, for logistics and information on crayfish trapping; Tony Rees and members of Merthyr Tydfil Angling Club water sample collection; Hampshire and Isle of Wight Wildlife Trust (Dr. Ben Rushbrook) and Environment Agency (Kerry Walsh, Emma McSwan, Kathy Friend) for assistance with sample collection and expertise in the Itchen and Medway and also Carlos Garcia de Leaniz for valuable advice on experimental design and comments to the final manuscript.

## Author contributions & competing interests

SC & CVR designed the study; CVR & TUW performed the analyses; JC & JJ contributed samples and information; SC & CVR wrote the paper with the help of all the authors. Authors declare that they have no competing interests.

## Data Accessibility

All data is currently included in the supplementary material and will be stored in Dryad upon acceptance if requested.

## REFERENCES

Agersnap, S., Larsen, W.B., Knudsen, S.W., Strand, D., Thomsen, P.F., Hesselsøe, M., Mortensen, P.B., Vrålstad, T., Møller, P.R., 2017. Monitoring of noble, signal and narrow-clawed crayfish using environmental DNA from freshwater samples. PLOS ONE 12, e0179261.

Alderman, D., 1996. Geographical spread of bacterial and fungal diseases of crustaceans. Revue scientifique et technique (International Office of Epizootics) 15, 603–632.

Alderman, D., Holdich, D., Reeve, I., 1990. Signal crayfish as vectors in crayfish plague in Britain. Aquaculture 86, 3–6.

Altizer, S., Harvell, D., Friedle, E., 2003. Rapid evolutionary dynamics and disease threats to biodiversity. Trends in Ecology and Evolution 18, 589–596.

Andreou, D., Arkush, K., Guégan, J., Gozlan, R., 2012. Introduced pathogens and native freshwater biodiversity: a case study of Sphaerothecum destruens. PLOS ONE 7.

Barnes, M., Turner, C., 2015. The ecology of environmental DNA and implications for conservation genetics Conservation Genetics.

Barnes, M.A., Turner, C.R., Jerde, C.L., Renshaw, M.A., Chadderton, W.L., Lodge, D.M., 2014. Environmental conditions influence eDNA persistence in aquatic systems. Environmental Science & Technology 48, 1819–1827.

Biggs, J., Ewald, N., Valentini, A., Gaboriaud, C., Dejean, T., Griffiths, R.A., Foster, J., Wilkinson, J.W., Arnell, A., Brotherton, P., Williams, P., Dunn, F., 2015. Using eDNA to develop a national citizen science-based monitoring programme for the great crested newt (Triturus cristatus). Biological Conservation 183, 19–28.

Bohmann, K., Evans, A., Gilbert, M.T.P., Carvalho, G.R., Creer, S., Knapp, M., Douglas, W.Y., de Bruyn, M., 2014. Environmental DNA for wildlife biology and biodiversity monitoring. Trends in Ecology & Evolution 29, 358–367.

Bubb, D.H., Thom, T.J., Lucas, M.C., 2005. The within-catchment invasion of the non-indigenous signal crayfish Pacifastacus leniusculus (Dana), in upland rivers. KMAE 376, 665–673.

Chapple, D.G., Simmonds, S.M., Wong, B.B., 2012. Can behavioral and personality traits influence the success of unintentional species introductions? Trends in Ecology & Evolution 27, 57–64.

Cooper, J., 2011. Anesthesia, analgesia and euthanasia of invertebrates. ILAR 52, 196–204.

Crowl, T.A., Crist, T.O., Parmenter, R.R., Belovsky, G., Lugo, A.E., 2008. The spread of invasive species and infectious disease as drivers of ecosystem change. Frontiers in Ecology and the Environment 6, 238–246.

Darling, J.A., Mahon, A.R., 2011. From molecules to management: adopting DNA-based methods for monitoring biological invasions in aquatic environments. Environmental Research 111, 978–988.

DEFRA, 2015. Common standards monitoring guidance for freshwater fauna, ed. JNCC.

Dejean, T., Valentini, A., Duparc, A., Pellier-Cuit, S., Pompanon, F., Taberlet, P., Miaud, C., 2011. Persistence of Environmental DNA in Freshwater Ecosystems. PLOS ONE 6, e23398.

Dejean, T., Valentini, A., Miquel, C., Taberlet, P., Bellemain, E., Miaud, C., 2012. Improved detection of an alien invasive species through environmental DNA barcoding: the example of the American bullfrog Lithobates catesbeianus. Journal of Applied Ecology 49, 953–959.

Díaz-Ferguson, E., Moyer, G., 2014. History, applications, methodological issues and perspectives for the use of environmental DNA (eDNA) in marine and freshwater environments. International Journal of Tropical Biology 62, 1273–1284.

Diéguez-Uribeondo, J., Huang, T., Cerenius, L., Söderhäll, K., 1995. Physiological adaptation of an Aphanomyces astaci strain isolated from the freshwater crayfish Procambarus clarkii. Mycological Research 99, 574–578.

Dougherty, M., Larson, E., Renshaw, M., Gantz, C., Egan, S., Erickson, D., Lodge, D., 2016. Environmental DNA (eDNA) detects the invasive rustycrayfish Orconectes rusticus at low abundances. Journal of Applied Ecology 53, 722–732.

Dunn, J., McClymont, H., Christmas, M., Dunn, A., 2009. Competition and parasitism in the native White Clawed Crayfish Austropotamobius pallipes and the invasive Signal Crayfish Pacifastacus leniusculus in the UK. Biological Invasions 11, 315–324.

Edgerton, B.F., Henttonen, P., Jussila, J., Mannonen, A., Paasonen, P., Taugbøl, T., Edsman, L., Souty-Grosset, C., 2004. Understanding the causes of disease in European freshwater crayfish. Conservation Biology 18, 1466–1474.

Eichmiller, J., Best, S., Sorensen, P., 2016. Effects of temperature and trophic state on degradation of environmental DNA in lake water. Environmental Science & Technology 50, 1859–1867.

Ficetola, G.F., Miaud, C., Pompanon, F., Taberlet, P., 2008. Species detection using environmental DNA from water samples. Biology Letters 4, 423–425.

Freeman, M., Turnball, J., Yeomans, W., Bean, C., 2010. Prospects for management strategies of invasive crayfish populations with an emphasis on biological control. Aquatic Conservation: Marine and Freshwater Ecosystems 20, 211–223.

Ganoza, C.A., Matthias, M.A., Collins-Richards, D., Brouwer, K.C., Cunningham, C.B., Segura, E.R., Gilman, R.H., Gotuzzo, E., Vinetz, J.M., 2006. Determining risk for devere Leptospirosis by molecular analysis of environmental surface waters for pathogenic Leptospira. PLOS Medicine 3, e308.

Gherardi, F., Aquiloni, L., Diéguez-Uribeondo, J., Tricarico, E., 2011. Managing invasive crayfish: is there a hope? Aquatic Sciences 73, 185–200.

Goldberg, C.S., Turner, C.R., Deiner, K., Klymus, K.E., Thomsen, P.F., Murphy, M.A., Spear, S.F., McKee, A., Oyler-McCance, S.J., Cornman, R.S., 2016. Critical considerations for the application of environmental DNA methods to detect aquatic species. Methods in Ecology and Evolution.

Griffiths, S.W., Collen, P., Armstrong, J.D., 2004. Competition for shelter among over-wintering signal crayfish and juvenile Atlantic salmon. Journal of Fish Biology 65, 436–447.

Guy, R.A., Payment, P., Krull, U.J., Horgen, P.A., 2003. Real-time PCR for quantification of Giardia and Cryptosporidium in environmental water samples and sewage. Applied and Environmental Microbiology 69, 5178–5185.

Haddaway, N.R., Wilcox, R.H., Heptonstall, R.E.A., Griffiths, H.M., Mortimer, R.J.G., Christmas, M., Dunn, A.M., 2012. Predatory functional response and prey choice identify predation differences between native/invasive and parasitised/unparasitised crayfish. PLOS ONE 7, e32229.

Holdich D.M., Lowery R.S., 1988. Pacifastacus leniusculus in North America and Europe, with details of the distribution of introduced and native crayfish species in Europe. In: Freshwater crayfish: Biology, management and exploitation (eds Holdich, D.M., Lowery, R.S.), pp.283–308. Croom Helm, London.

Holdich, D.M., Reynolds, J.D., Souty-Grosset, C., Sibley, P.J., 2009. A review of the ever increasing threat to European crayfish from non-indigenous crayfish species. Knowledge and Management of Aquatic Ecosystems, 11.

Ikeda, K., Doi, H., Tanaka, K., Kawai, T., Negishi, J., 2016. Using environmental DNA to detect an endangered crayfish Cambaroides japonicus in streams. Conservation Genetic Resources 8, 231–234.

IUCN, 2017. IUCN Red List of Threatened Species: Austropotamobius pallipes.

James, J., Nutbeam-Tuffs, S., Cable, J., Mrugała, A., Viñuela-Rodriguez, N., Petrusek, A., Oidtmann, B., 2017. The prevalence of Aphanomyces astaci in invasive signal crayfish from the UK and implications for native crayfish conservation. Parasitology 144, 1–8.

James, J., Slater, F., Cable, J., 2014a. A.L.I.E.N. databases: addressing the lack in establishment of non-natives databases. Crustaceana 87, 1192–1199.

James, J., Slater, F., Vaughan, I., Young, K., Cable, J., 2014b. Comparing the ecological impacts of native and invasive crayfish: could native species’ translocation do more harm than good? Oecologia 178. 309–316..

Jane, S.F., Wilcox, T.M., McKelvey, K.S., Young, M.K., Schwartz, M.K., Lowe, W.H., Letcher, B.H., Whiteley, A.R., 2015. Distance, flow and PCR inhibition: eDNA dynamics in two headwater streams. Molecular Ecology Resources 15, 216–227.

Jeschke, J.M., Bacher, S., Blackburn, T.M., Dick, J.T., Essl, F., Evans, T., Gaertner, M., Hulme, P.E., Kühn, I., MrugałA, A., 2014. Defining the impact of non-native species. Conservation Biology 28, 1188–1194.

Kelly, R.P., Port, J.A., Yamahara, K.M., Martone, R.G., Lowell, N., Thomsen, P.F., Mach, M.E., Bennett, M., Prahler, E., Caldwell, M.R., Crowder, L.B., 2014. Harnessing DNA to improve environmental management. Science 344, 1455–1456.

Kirjavainen, J., Sipponen, M., 2004. Environmental benefit of different crayfish management strategies in Finland. Fisheries Management and Ecology 11, 213–218.

Klymus, K.E., Richter, C.A., Chapman, D.C., Paukert, C., 2015. Quantification of eDNA shedding rates from invasive bighead carp Hypophthalmichthys nobilis and silver carp Hypophthalmichthys molitrix. Biological Conservation 183, 77–84.

Kozubíková, E., Filipová, L., Kozák, P., Ďuriš, Z., Martín, M.P., Diéguez-Urbibeondo, J., Oidtmann, B., Petrusek, A., 2008. Prevalence of crayfish plague pathogen Aphanomyces astaci in invasive American crayfishes in the Czech Republic. Conservation Biology 23, 1204–1213.

Laramie, M.B., Pilliod, D.S., Goldberg, C.S., 2015. Characterizing the distribution of an endangered salmonid using environmental DNA analysis. Biological Conservation 183, 29–37.

Lodge, D.M., Deines, A., Gherardi, F., Yeo, D.C., Arcella, T., Baldridge, A.K., Barnes, M.A., Chadderton, W.L., Feder, J.L., Gantz, C.A., 2012. Global introductions of crayfishes: evaluating the impact of species invasions on ecosystem services. Annual Review of Ecology, Evolution, and Systematics 43, 449–472.

Lodge, D.M., Simonin, P.W., Burgiel, S.W., Keller, R.P., Bossenbroek, J.M., Jerde, C.L., Kramer, A.M., Rutherford, E.S., Barnes, M.A., Wittmann, M.E., Chadderton, W.L., Apriesnig, J.L., Beletsky, D., Cooke, R.M., Drake, J.M., Egan, S.P., Finnoff, D.C., Gantz, C.A., Grey, E.K., Hoff, M.H., Howeth, J.G., Jensen, R.A., Larson, E.R., Mandrak, N.E., Mason, D.M., Martinez, F.A., Newcomb, T.J., Rothlisberger, J.D., Tucker, A.J., Warziniack, T.W., Zhang, H., 2016. Risk analysis and bioeconomics of invasive species to inform policy and management. Annual Review of Environment and Resources 41, 453–488.

Lymbery, A., Morine, M., Kanani, H., Beatty, S., Morgan, D., 2014. Co-invaders: The effects of alien parasites on native hosts. International Journal for Parasitology: Parasites and Wildlife 3, 171–177.

Mauvisseau, Q., Coignet, A., Delaunay, C., Pinet, F., Bouchon, D., Souty-Grosset, C., 2018. Environmental DNA as in efficient tool for detecting invasive crayfishes in freshwater ponds. Hydrobiologia 805, 163–175.

Miaud, C., Dejean, T., Savard, K., Millery-Vigues, A., Valentini, A., Gaudin, N.C.G., Garner, T.W., 2016. Invasive North American bullfrogs transmit lethal fungus Batrachochytrium dendrobatidis. Biological Invasions 18, 2299–2308.

Moorhouse, T., Macdonald, D., 2015. Are invasions worse in freshwater then terrestrial ecosystems? Wiley Interdisciplinary Reviews: Water 2, 1–8.

Moyer, G., Díaz-Ferguson, E., Hill, J., Shea, C., 2014. Assessing Environmental DNA detection in controlled lentic systems. PLOS ONE 9, 1–9.

NBN, 2009. Crayfish (Crustacea; Astacura) data for Britain and Ireland to 2003, Oxfordshire.

Olden, J.D., Kennard, M.J., Lawler, J.J., Leroy, P.N., 2010. Challenges and opportunities in implementing managed relocation for conservation of freshwater species. Conservation Biology 25, 40–47.

Olson, Z.H., Briggler, J.T., Williams, R.N., 2012. An eDNA approach to detect eastern hellbenders (Cryptobranchus a. alleganiensis) using samples of water. Wildlife Research 39, 629–636.

Peay, S., 2009. Invasive non-indigenous crayfish species in Europe: Recommendations on managing them. Knowledge and Management of Aquatic Ecosystems 3, 394–395.

Peeler, E.J., Oidtmann, B.C., Midtlyng, P.J., Miossec, L., Gozlan, R.E., 2011. Non-native aquatic animals introductions have driven disease emergence in Europe. Biological Invasions 13, 1291–1303.

Peters, J.A., Lodge, D.M., 2013. Habitat, predation, and coexistence between invasive and native crayfishes: prioritizing lakes for invasion prevention. Biological Invasions 15, 2489–2502.

Pilliod, D.S., Goldberg, C.S., Arkle, R.S., Waits, L.P., 2013. Estimating occupancy and abundance of stream amphibians using environmental DNA from filtered water samples. Canadian Journal of Fisheries and Aquatic Sciences 70, 1123–1130.

Pilliod, D.S., Goldberg, C.S., Arkle, R.S., Waits, L.P., 2014. Factors influencing detection of eDNA from a stream-dwelling amphibian. Molecular Ecology Resources 14, 109–116.

Pintor, L.M., Sih, A., Bauer, M.L., 2008. Differences in aggression, activity and boldness between native and introduced populations of an invasive crayfish. Oikos 117, 1629–1636.

Pochon, X., Bott, N.J., Smith, K.F., Wood, S.A., 2013. Evaluating detection limits of next-generation sequencing for the surveillance and monitoring of international marine pests. PLOS ONE 8, e73935.

Ramón-Laca, A., Gleeson, D., Yockney, I., Perry, M., Nugent, G., Forsyth, D.M., 2014. Reliable discrimination of 10 ungulate species using high resolution melting analysis of faecal DNA. PLOS ONE 9, e92043.

Randolph, S.E., Rogers, D.J., 2010. The arrival, establishment and spread of exotic diseases: patterns and predictions. Nature Reviews in Microbiology 8, 361–371.

Rees, H., Maddison, B., Middleditch, D., Patmore, J., Gough, K., 2014. The detection of aquatic animal species using environmental DNA – a review of eDNA as a survey tool in ecology. Journal of Applied Ecology 51, 1450–1459.

Ririe, K.M., Rasmussen, R.P., Wittwer, C.T., 1997. Product differentiation by analysis of DNA melting curves during the polymerase chain reaction. Analytical biochemistry 245, 154–160.

Rushbrook, B., 2014. Crayfish Conservation in Hampshire’s Chalk Streams.

Russell, J.C., Blackburn, T.M., 2017. The rise of invasive species denialism. Trends in Ecology & Evolution 32, 3–6.

Sandodden, R., Johnsen, S., 2010. Eradication of introduced signal crayfish Pasifastacus leniusculus using the pharmaceutical BETAMAX VET.^®^. Aquatic Invasions 5, 75–81.

Schrimpf, A., Pârvulescu, L., Copilas-Ciocianu, D., Petrusek, A., Schulz, R., 2012. Crayfish plague pathogen detected in the Danube Delta-a potential threat to freshwater biodiversity in southeastern Europe. Aquatic Invasions 7, 503–510.

Sigsgaard, E.E., Carl, H., Møller, P.R., Thomsen, P.F., 2015. Monitoring the near-extinct European weather loach in Denmark based on environmental DNA from water samples. Biological Conservation 183, 46–52.

Simberloff, D., Martin, J.-L., Genovesi, P., Maris, V., Wardle, D.A., Aronson, J., Courchamp, F., Galil, B., García-Berthou, E., Pascal, M., Pyšek, P., Sousa, R., Tabacchi, E., Vilà, M., 2013. Impacts of biological invasions: what’s what and the way forward. Trends in Ecology & Evolution 28, 58–66.

Smart, A., Tingley, R., Weeks, A., van Rooyen, A., McCarthy, M., 2015. Environmental DNA sampling is more sensitive than a traditional survey technique for detecting an aquatic invader. Ecological Applications 25, 1944–1952.

Spear, S.F., Groves, J.D., Williams, L.A., Waits, L.P., 2015. Using environmental DNA methods to improve detectability in a hellbender (Cryptobranchus alleganiensis) monitoring program. Biological Conservation 183, 38–45.

Strand, D.A., Jussila, J., Johnsen, S.I., Viljamaa-Dirks, S., Edsman, L., Wiik-Nielsen, J., Viljugrein, H., Engdahl, F., Vrålstad, T., 2014. Detection of crayfish plague spores in large freshwater systems. Journal of Applied Ecology 51, 544–553.

Strauss, A., White, A., Boots, M., 2012. Invading with biological weapons: the importance of disease-mediated invasions. Functional Ecology 26, 1249–1261.

Taberlet, P., Coissac, E., Hajibabaei, M., Rieseberg, L.H., 2012. Environmental DNA. Molecular Ecology 21, 1789–1793.

Takahara, T., Minamoto, T., Doi, H., 2013. Using Environmental DNA to estimate the distribution of an invasive fish species in ponds. PLoS ONE 8, e56584.

Thomsen, P., Kielgast, J., Iversen, L.L., Wiuf, C., Rasmussen, M., Gilbert, M.T.P., Orlando, L., Willerslev, E., 2012. Monitoring endangered freshwater biodiversity using environmental DNA. Molecular Ecology 21, 2565–2573.

Thomsen, P.F., Willerslev, E., 2015. Environmental DNA – An emerging tool in conservation for monitoring past and present biodiversity. Biological Conservation 183, 4–18.

Tréguier, A., Paillisson, J.-M., Dejean, T., Valentini, A., Schlaepfer, M., Roussel, J.-M., 2014. Environmental DNA surveillance for invertebratespecies: advantages and technical limitations todetect invasive crayfish Procambarus clarkii infreshwater ponds. Journal of Applied Ecology 51, 871–879.

Turner, C.R., Uy, K.L., Everhart, R.C., 2015. Fish environmental DNA is more concentrated in aquatic sediments than surface water. Biological Conservation 183, 93–102.

Unestam, T., 1966. Studies on the crayfish plague fungus Aphanomyces astaci. Physiologia Plantarum 19, 1110–1119.

Vander Zanden, M.J., Hansen, G.J., Higgins, S.N., Kornis, M.S., 2010. A pound of prevention, plus a pound of cure: early detection and eradication of invasive species in the Laurentian Great Lakes. Journal of Great Lakes Research 36, 199–205.

Vilcinskas, A., 2015. Pathogens as biological weapons of invasive species. PLoS Pathogens 11.

Vitousek, P.M., D Antonio, C.M., Loope, L.L., Westbrooks, R., 1996. Biological invasions as global environmental change. American Scientist 84, 468.

Vrålstad, T., Knutsen, A., Tengs, T., Holst-Jensen, A., 2009. A quantitative TaqMan MGB real-time polymerase chain reaction based assay for detection of the causative agent of crayfish plague Aphanomyces astaci. Veterinary Microbiology 137, 146–155.

Westman, K., Savolainen, R., Julkunen, M., 2002. Replacement of the native crayfish Astacus astacus by the introduced species Pacifastacus leniusculus in a small, enclosed Finnish lake: a 30-year study. Ecography 25, 53–73.

Wilson, C.C., Wozney, K.M., Smith, C.M., 2016. Recognizing false positives: synthetic oligonucleotide controls for environmental DNA surveillance. Methods in Ecology and Evolution 7, 23–29.

Wittwer, C., Stoll, S., Strand, D., Vrålstad, T., Nowak, C., Thines, M., 2018. eDNA-based crayfish plague monitoring is superior to conventional trap-based assessments in year-round detection probability. Hydrobiologia 807, 87–97.

Wye & Usk Foundation, 2012. The Tubney Charitable Trust.

Yang, S., Ramachandran, P., Rothman, R., Hsieh, Y.-H., Hardick, A., Won, H., Kecojevic, A., Jackman, J., Gaydos, C., 2009. Rapid identification of biothreat and other clinically relevant bacterial species by use of Universal PCR coupled with High-Resolution Melting Analysis. Journal of Clinical Microbiology 47, 2252–2255.

Ye, J., McGinnis, S., Madden, T.L., 2006. BLAST: improvements for better sequence analysis. Nucleic Acids Research 34, W6–W9.

Zaiko, A., Minchin, D., Olenin, S., 2014. “The day after tomorrow”: anatomy of an ‘r’strategist aquatic invasion. Aquatic Invasions 9, 145–155.

